# Metabolic inequality in microbial communities

**DOI:** 10.64898/2026.04.14.718602

**Authors:** Emmi A. Mueller, Jay T. Lennon

## Abstract

How metabolic activity is partitioned among individuals determines the scaling of cellular physiology to higher levels of biological organization. Yet the mechanisms that generate this heterogeneity and shape its distribution remain largely unresolved. We quantified single-cell metabolism in microbial communities spanning aquatic, terrestrial, and host-associated ecosystems. Across more than one million cells, metabolic activity followed a long-tailed distribution best described by a lognormal model, with a small subset of individuals contributing disproportionately to community metabolism. In some cases, the most active 20% of cells accounted for over 90% of metabolic output, but this pattern became less pronounced in more productive environments. To assess the consequences of metabolic inequality, we developed a stochastic simulation model linking single-cell activity to community respiration via enzyme kinetics. Because respiration responds nonlinearly to enzyme activity, variation among cells does not translate proportionally into ecosystem-level fluxes. As a result, ignoring metabolic heterogeneity can bias estimates of community respiration by up to 60%. Our findings reveal a general pattern of metabolic inequality in microbial communities that holds across a wide range of habitats. Accounting for this structure is critical for understanding how microorganisms regulate ecosystem processes and for improving predictions of large-scale biogeochemical dynamics.

**Significance:** Inequality and disparity are common features of social, economic, and physical systems. Such patterns also arise in nature, where a small fraction of individuals accounts for an outsized share of biological output, including reproduction, immunity, and diversity. Here, we show that metabolic activity in microbial communities follows a characteristic right-skewed distribution across diverse ecosystems, including lakes, soils, ocean plankton, marine sediments, and mammalian guts. Instead of the rich getting richer, metabolism becomes more evenly distributed among cells in more productive environments. By explicitly representing metabolic inequality, we can reduce bias in estimates of microbial activity and improve the scaling of cellular processes to ecosystem-level fluxes.

---

Metabolism is the sum of life’s chemical reactions and forms a unifying foundation for ecological theory, linking structure and function across all scales and levels of biological organization [1–4]. While metabolic rates vary widely among taxa and constrain ecological processes [5–8], substantial variation also occurs within species. Metabolic rates are highly plastic, responding to abiotic and biotic conditions that fluctuate across space and time [9–11]. Even in the absence of genetic or environmental differences, stochastic processes can introduce variability that influences the metabolic output of a population [12, 13].

Among living systems, microorganisms are particularly well-suited for examining individual-level metabolic variation. They encode a vast repertoire of catalytic functions [14–16], but their contributions to ecosystem processes depend not only on which taxa are present but also on physiological differences among individual cells [17, 18]. This variation challenges mean-field assumptions and makes microbial communities ideal for investigating how individual traits scale to influence collective outcomes. For example, many metabolic tasks rely on cooperative interactions that no single cell can perform alone [19]. Such interactions give rise to metabolic networks that enable microbes to thrive in diverse habitats and drive transformations essential to the stability of ecosystems, from host microbiomes to the biosphere [20, 21]. As a result, microbial traits and physiological properties are increasingly being integrated into models that aim to improve predictions of Earth system processes [22–26]. Yet these models often overlook the causes and consequences of metabolic variation among individual cells, reducing ecological dynamics to averages that mask underlying heterogeneity [27].

Without understanding how metabolic activity is distributed among individuals, simplifications are made that can compromise system-level predictions. Many models assume that all individuals contribute equally, yielding a uniform distribution in which each of the *N* individuals accounts for 1*/N* of the total activity (*A*_*total*_) [28, 29]. Even when variation in metabolism is explicitly represented, for instance using a Gaussian distribution, the underlying assumption remains one of symmetry around a mean value (*A*_*total*_*/N* ).

However, complex assemblages of microorganisms are unlikely to exhibit such equality. Most cells do not operate at their full metabolic potential, and many can persist in an inactive state for extended periods of time [30]. If cells effectively exist in on/off states, activity may follow a bimodal distribution. If instead activity spans a physiological continuum, long-tailed distributions may emerge, consistent with the Pareto principle, in which a small fraction of individuals accounts for a disproportionately large share of total output [31]. Skewed distributions such as the lognormal, gamma, or power-law can arise from additive, multiplicative, or stochastic processes that govern key metabolic functions, including gene expression, biomass accumulation, and variation in growth rate [32–35].

Determining whether microbial communities exhibit a characteristic distribution of activity is essential for developing a general theory of metabolic ecology, which seeks to explain how energy and metabolism structure biological organization across scales [2]. A model that captures both the central tendency and the degree of inequality in activity (Fig. 1) provides a basis for investigating factors that influence metabolic organization, including growth strategies, functional traits, and resource availability. Because productivity governs energy and resource supply, it likely influences the distribution of metabolic activity within communities [8, 36]. For example, if productivity disproportionately enhances access to resources for already active individuals, metabolic distributions may become more skewed, consistent with preferential attachment dynamics [37, 38]. Alternatively, if productivity broadens access to resources among individuals, activity may become more evenly distributed [39]. Taken together, shifts in the shape of metabolic distributions along environmental gradients can reveal underlying constraints and help identify general principles governing microbial metabolism.

**Figure 1.**
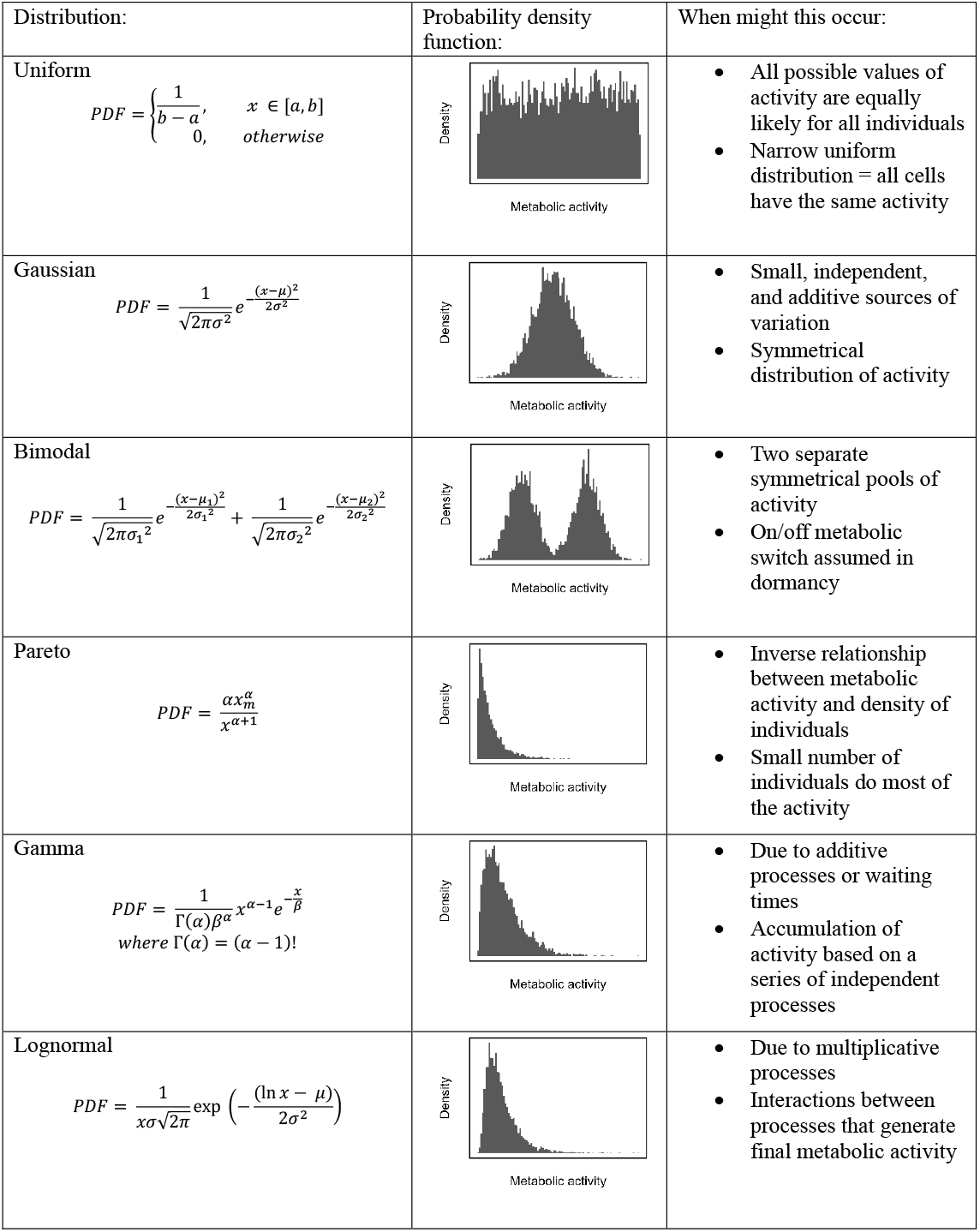
Metabolic distributions. Potential distributions of metabolic activity at the single-cell level and the mechanisms that generate them. Each row shows the probability density function (PDF), its visualization, and a summary of potential mechanisms and their implications for community-level metabolic activity.

In this study, we determined how metabolism is distributed among individuals within species and across microbial communities in diverse ecosystems. Using flow cytometry, we quantified reductase activity at the single-cell level and used it as a proxy for redox activity. Applying this approach, we tested how metabolic activity distributions vary with growth regimes and productivity and evaluated the generality of these patterns across diverse microbiomes. Finally, we developed a stochastic simulation model to evaluate the consequences of metabolic inequality for ecosystem processes. Together, these approaches provide a framework for linking metabolic organization to ecological theory and improving predictions of biogeochemical dynamics.

## Results and Discussion

Across a wide range of growth conditions and ecosystems, we observed a recurring pattern of metabolic inequality within microbial communities. Many cells exhibited low metabolic activity, while relatively few exhibited high activity, resulting in consistently right-skewed distributions best described by a lognormal model. This pattern was conserved in a diverse collection of bacterial isolates, suggesting that the community-level distribution does not arise simply from combining distinct species-level distributions but instead reflects a more general feature of microbial physiology. We next asked how environmental conditions shape this inequality. Because productivity governs resource availability, it likely influences how metabolic activity is partitioned among individuals. Rather than amplifying disparities, metabolism was more evenly distributed among individuals in productive environments. Applying the principles of nonlinear averaging, we show that accounting for skewed activity distributions alters estimates of aggregate metabolism and can improve predictions of ecosystem-scale dynamics.

### 1. Metabolic activity is unequally distributed

Uneven distributions of activity are more likely than uniform ones. There are simply many more ways for activity to be unevenly distributed among individuals [40]. As a result, heterogeneity emerges as the most probable outcome, even in the absence of specific physiological or ecological drivers. In many complex systems, this tendency gives rise to inequality, in which a small fraction of individuals accounts for most of the total output. Such uneven distributions may follow the Pareto principle, where roughly 80% of output is generated by 20% of producers. To test whether microbial communities exhibit these patterns, we quantified the fraction of activity attributed to the top 20% of cells across three contrasting growth regimes: (1) batch cultures, where resource depletion, biomass production, and waste accumulation drive transitions through lag, exponential, and stationary phases; (2) continuous cultures (chemostats), where dilution rate determines growth at steady state through the balance of resource supply and washout; and (3) environmental samples from a freshwater reservoir, where a hydrological flow path establishes a gradient of productivity.

Metabolic inequality was widespread across growth regimes, with the top 20% of individuals contributing over 90% of total activity in some cases (Fig. 2). Batch culture exhibited the greatest variability (Fig. 3). During lag phase, when resources were replete, about 75% of total activity was produced by the 20% most active cells (Fig. 3B, S1, & S2). During exponential and stationary phases, activity became more evenly distributed (32–41%), yet the most active individuals still contributed about 1.5 times their expected share of total activity (Fig. 3C-D, S1, & S2). In contrast, environmental samples and continuous cultures exhibited narrower ranges, with 51–90% and 66–91% of total metabolism attributable to the most active cells, respectively (Fig. 2). Even under optimal conditions, a substantial fraction of cells remained in a state of reduced metabolic activity, consistent with bet-hedging strategies in which some individuals sacrifice instantaneous fitness to buffer populations against environmental change [41–43].

**Figure 2.**
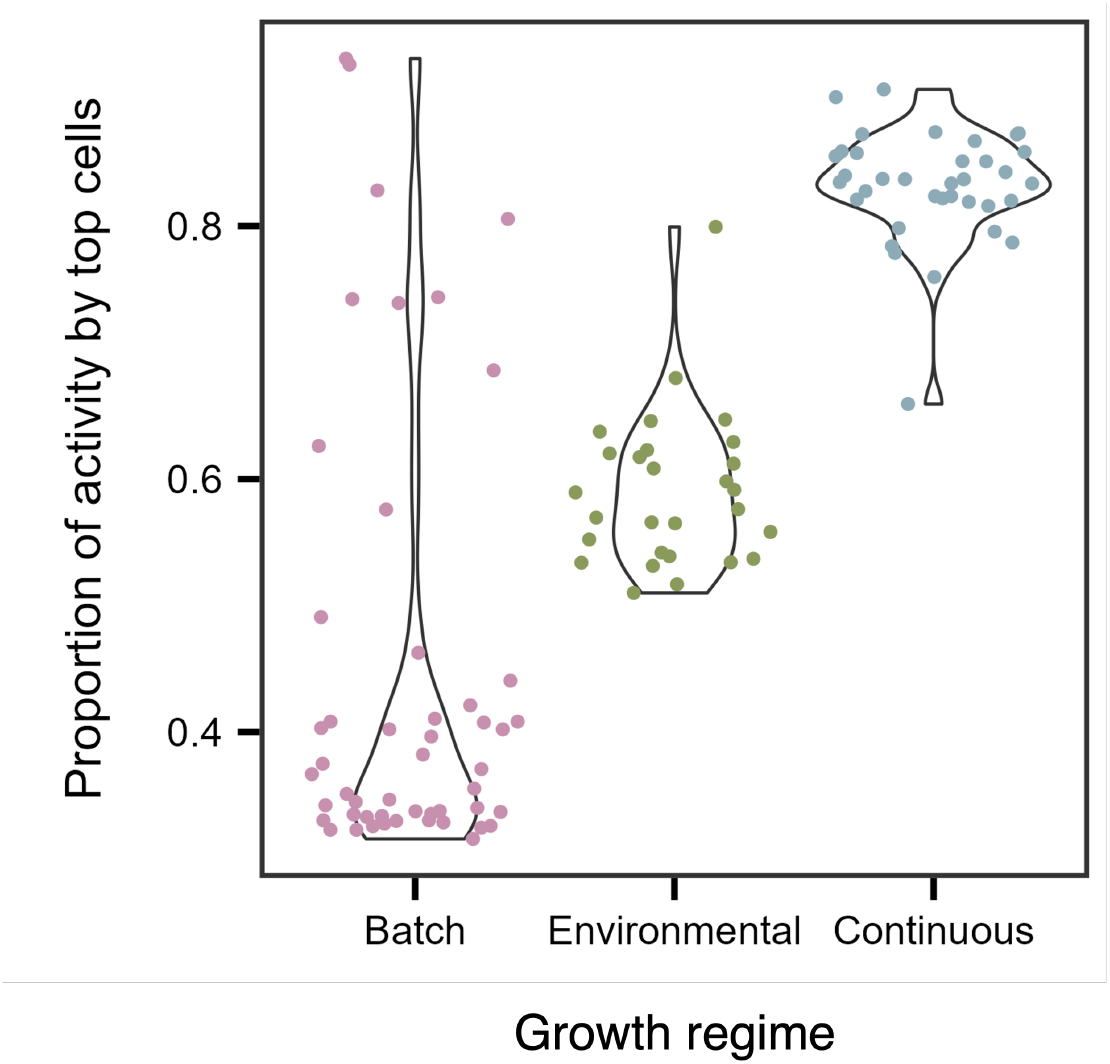
Metabolic activity is unevenly distributed. Violin plots showing that the 20% most active cells contribute a large but variable fraction of total community-level activity. Batch cultures exhibit the widest range, consistent with shifts in metabolic distributions across growth phases (lag, exponential, and stationary). In contrast, environmental samples and continuous cultures (chemostats) show narrower ranges.

**Figure 3.**
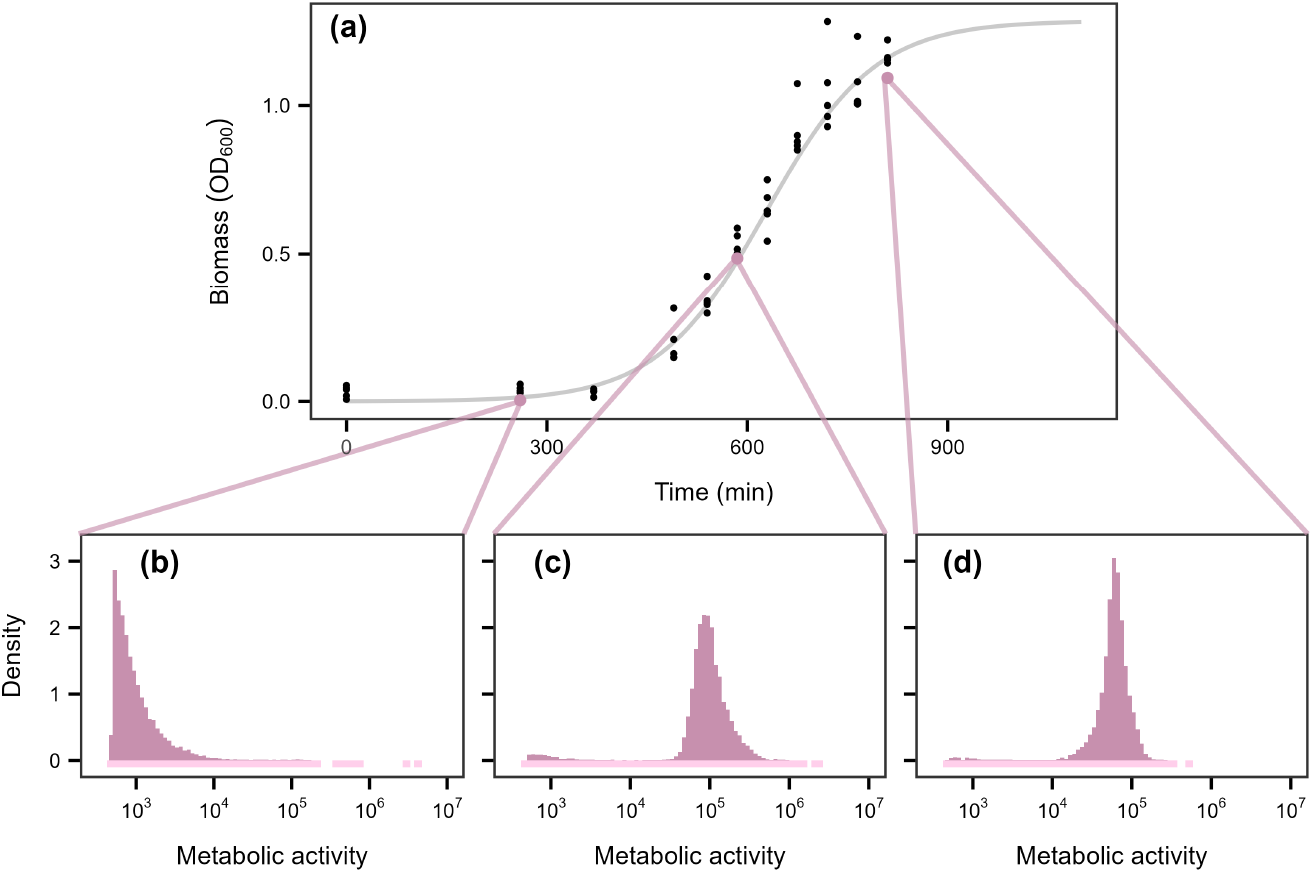
Growth-phase dependent shifts in metabolic activity. **(a)** Microbial biomass of a lake water community grown in batch culture over time. In the lower panels **(b-d)**, metabolic activity is reported as relative fluorescent units (RFU). Density plots show the distribution of metabolic activity across three growth stages: lag, exponential, and stationary. To highlight the tails of the distribution, low-density histogram bins are extended below zero using a lighter tint of the histogram color. **(b)** In lag phase, metabolic activity was low and highly right-skewed, with most cells exhibiting activity values below 10^5^ (median = 810; mean = 2,461). **(c)** In exponential phase, metabolic activity increased by approximately two orders of magnitude, and the distribution remained right-skewed (median = 93,083; mean = 111,569). **(d)** In stationary phase, the distribution became more even, with activity levels similar to those in exponential phase (median = 60,734; mean = 63,203).

### 2. A characteristic distribution for metabolic activity

Identifying how activity is distributed among individuals is essential for understanding the causes and consequences of phenotypic heterogeneity, advancing metabolic theory, and improving our broader understanding of functional ecology [44]. Yet, no widely accepted model captures how metabolism is partitioned or how resulting distributions shift across environmental gradients and ecosystem types. Parameterizing a distributional model provides a concise way to summarize complex datasets while also establishing a common framework for cross-study comparisons. Moreover, these models can serve as powerful tools for ruling out implausible explanations, generating testable hypotheses, and providing baselines for assessing how populations, communities, and ecosystems respond to perturbations.

Using over 500,000 single-cell reductase measurements, we evaluated whether a single distribution could reliably describe the metabolic activity of aquatic microorganisms under different growth regimes. Consistently, the lognormal distribution was the best-fitting model, outperforming all others (uniform, bimodal, Gaussian, gamma, and Pareto; see Fig. 1) based on both *r*^2^ values (Fig. 4A; Table S1 & S2; Batch: *F*_3_ = 84.91, *P <* 0.0001; Continuous: *F*_3_ = 449.4, *P <* 0.0001; Environmental: *F*_3_ = 1588, *P <* 0.0001) and Kolmogorov-Smirnov (K-S) statistics (Fig. 4B; Table S1 & S2; Batch: *F*_3_ = 55.2, *P <* 0.0001; Continuous: *F*_3_ = 803, *P <* 0.0001; Environmental: *F*_3_ = 1065, *P <* 0.0001). According to Akaike Information Criterion (AIC) and Bayesian Information Criterion (BIC), the lognormal provided the best fit for all continuous-culture and environmental samples tested (28 of 28 and 36 of 36, respectively). In batch cultures, the best-fitting models were more variable: the lognormal ranked best in 7 of 50 samples, compared to 16 for the gamma, 25 for the Gaussian, and 2 for the Pareto (Fig. S3). Bootstrapped 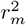 values and quantile–quantile (Q–Q) plots further supported the lognormal model, which achieved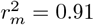 compared with 0.83 for the gamma 0.72 for the Gaussian, and 0.80 for the Pareto (Fig. 4C-F).

**Figure 4.**
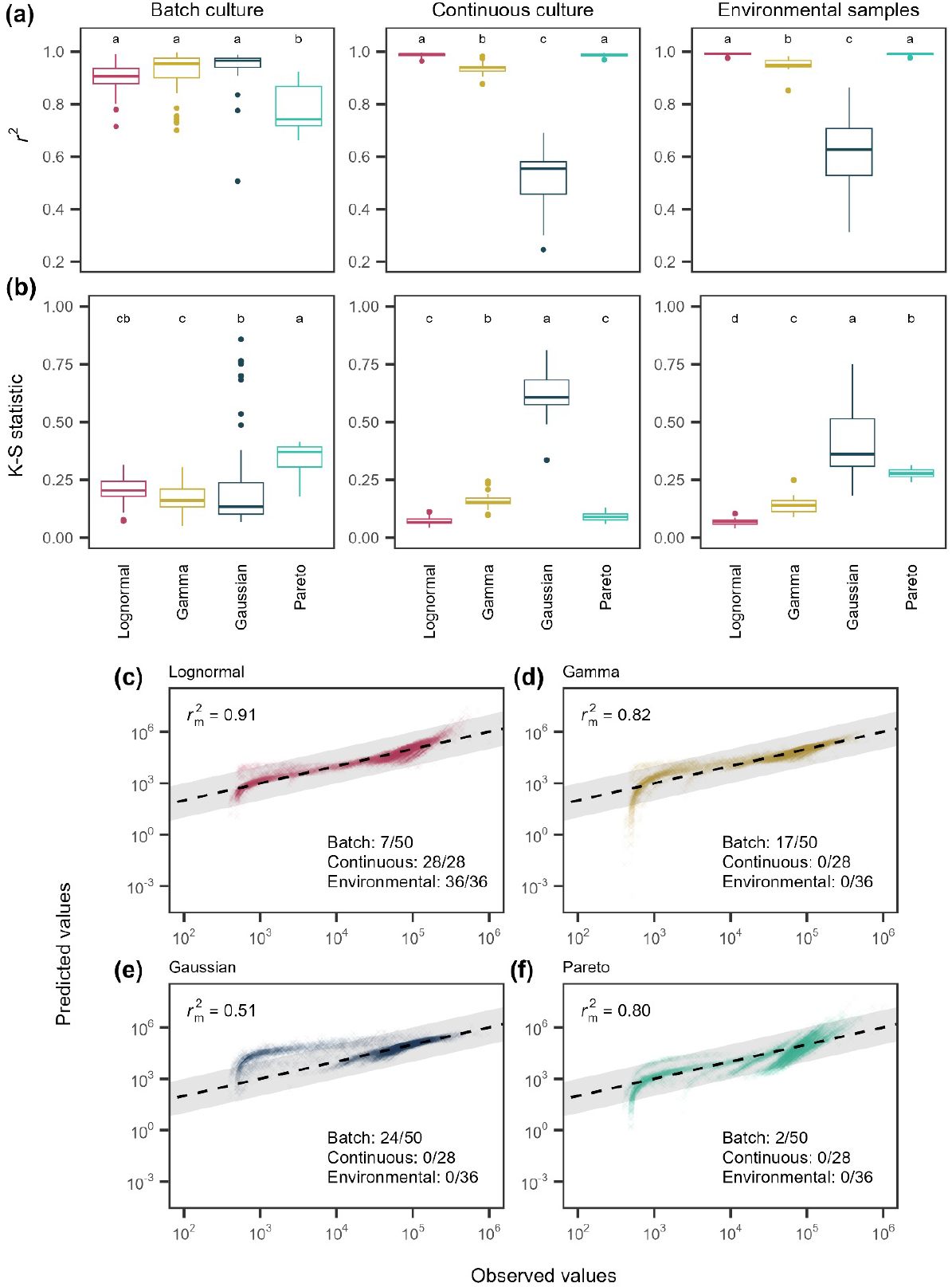
Model fits to single-cell metabolic activity distributions. Metabolic activity distributions for microbial communities across three growth regimes (batch culture, continuous culture, environmental samples) were fit with four candidate models: lognormal, gamma, Gaussian, and Pareto. **(A)** Coefficients of determination (*r*^2^) for each sample–model fit; letters denote groupings based on *post hoc* Tukey tests. **(B)** Kolmogorov–Smirnov (K–S) statistics, where lower values indicate better fit. **(C–F)** Quantile–quantile plots comparing observed and predicted activity for the lognormal, gamma, Gaussian, and Pareto models, respectively. Points represent 10,000 subsampled individuals across growth conditions. Solid lines indicate the 1:1 relationship, and shaded regions represent *±*1 order of magnitude in activity. Numbers indicate the number of samples for which each model was the best fit, based on ΔAIC and ΔBIC *>* 2. Across metrics and conditions, the lognormal model provides the best overall fit among candidate models.

The robustness of the lognormal model in describing reductase activity provides insight into the potential mechanisms underlying the pattern of microbial inequality. Specifically, it allowed us to eliminate some *a priori* models that fit the data poorly (Fig. 1). For example, the bimodal model failed to capture activity patterns, suggesting that metabolism in complex communities cannot be explained by simple on/off regulation, while the Gaussian model, which assumes additive, independent effects around a mean, also performed poorly. The gamma model, which would suggest that metabolism is the sum of sequential stochastic processes with exponential waiting times, provided a better fit by capturing right-skew in activity [45].

Among all models considered, the Pareto and lognormal distributions provided the best overall fit. The distinction between these distributions offers insight into the mechanisms generating metabolic inequality. While both capture right-skewed patterns, they differ in their interpretation. Pareto distributions are typically associated with unbounded reinforcing feedbacks that produce heavy-tailed distributions, whereas lognormal distributions are often associated with stochastic proportional changes around an equilibrium, generating finite variance and lighter tails [31]. The lognormal therefore represents a common outcome of such multiplicative dynamics, in which a small subset of cells contributes disproportionately to total metabolism [46, 47]. Its generality also explains why lognormal distributions commonly arise in diverse biological contexts, including transcript levels [48], body sizes [49], species abundances [50], and cell growth dynamics [35].

### 3. Species-level contributions to metabolic inequality

Ecological communities are composed of species with distinct life-history strategies and functional traits. As a result, the community-level distribution of metabolic activity must reflect its constituent taxa and may arise from the combination of distinct species-level distributions. To evaluate whether individual species share a common metabolic distribution, we measured single-cell reductase activity in a diverse set of heterotrophic bacteria grown under oligotrophic conditions. Across all bacterial isolates, metabolic activity followed a lognormal distribution (Fig. S4; Table S3 & S4). This shared form suggests that the community-level pattern does not arise from the pooling of fundamentally different species-level distributions. At the same time, the degree of metabolic inequality varied among species, even under identical growth conditions (Fig. S5). Most species had moderate skew, quantified by the log-scale standard deviation (*σ*) of the fitted lognormal distribution. A subset, however, exhibited pronounced inequality, with the most active 20% of cells accounting for more than 70% of total activity (Fig. S5). Notably, *σ* showed a strong phylogenetic signal consistent with Brownian motion (Pagel’s *λ* = 0.99, *P <* 0.0001), indicating that evolutionary divergence among species has the potential to shape community-level metabolic distributions.

### 4. Metabolism is more evenly distributed in productive environments

Productivity is the rate at which energy is converted into new biomass. It sets a fundamental constraint on the amount of biological work that can be performed and, as a result, shapes patterns of abundance, diversity, body size, species interactions, and ecosystem functioning [51]. Thus, we hypothesized that productivity should influence the distribution of metabolic activity among individuals within a community. On the one hand, increased productivity could disproportionately fuel already active individuals, thereby amplifying metabolic inequality. On the other hand, increased productivity could elevate activity across individuals while reducing differences among them, resulting in a more even metabolic distribution.

To test these contrasting scenarios, we examined the relationship between bacterial productivity (BP) and metabolic inequality across two complementary datasets. In a freshwater reservoir, where BP varied longitudinally along the hydrological flow path (Fig. S6; Table S5), metabolic skew (*σ*) declined with increasing productivity (Fig. 5). Similarly, in laboratory chemostats, experimental manipulation of BP in communities derived from the freshwater reservoir (Fig. **??**; Table S5) led to a decline in metabolic skew (Fig. 5 & S8). Using an indicator-variable multiple regression, we found that the baseline skew, represented by the intercept, was approximately twofold higher in chemostats than in the reservoir ecosystem. However, the slopes relating skew to log_10_(BP) were statistically indistinguishable (-0.14 ± 0.031), indicating that metabolic activity becomes more evenly distributed with increasing productivity in aquatic microbial communities (*R*^2^ = 0.96, *F*_3,60_ = 488.5, *P <* 0.0001, Table S6).

**Figure 5.**
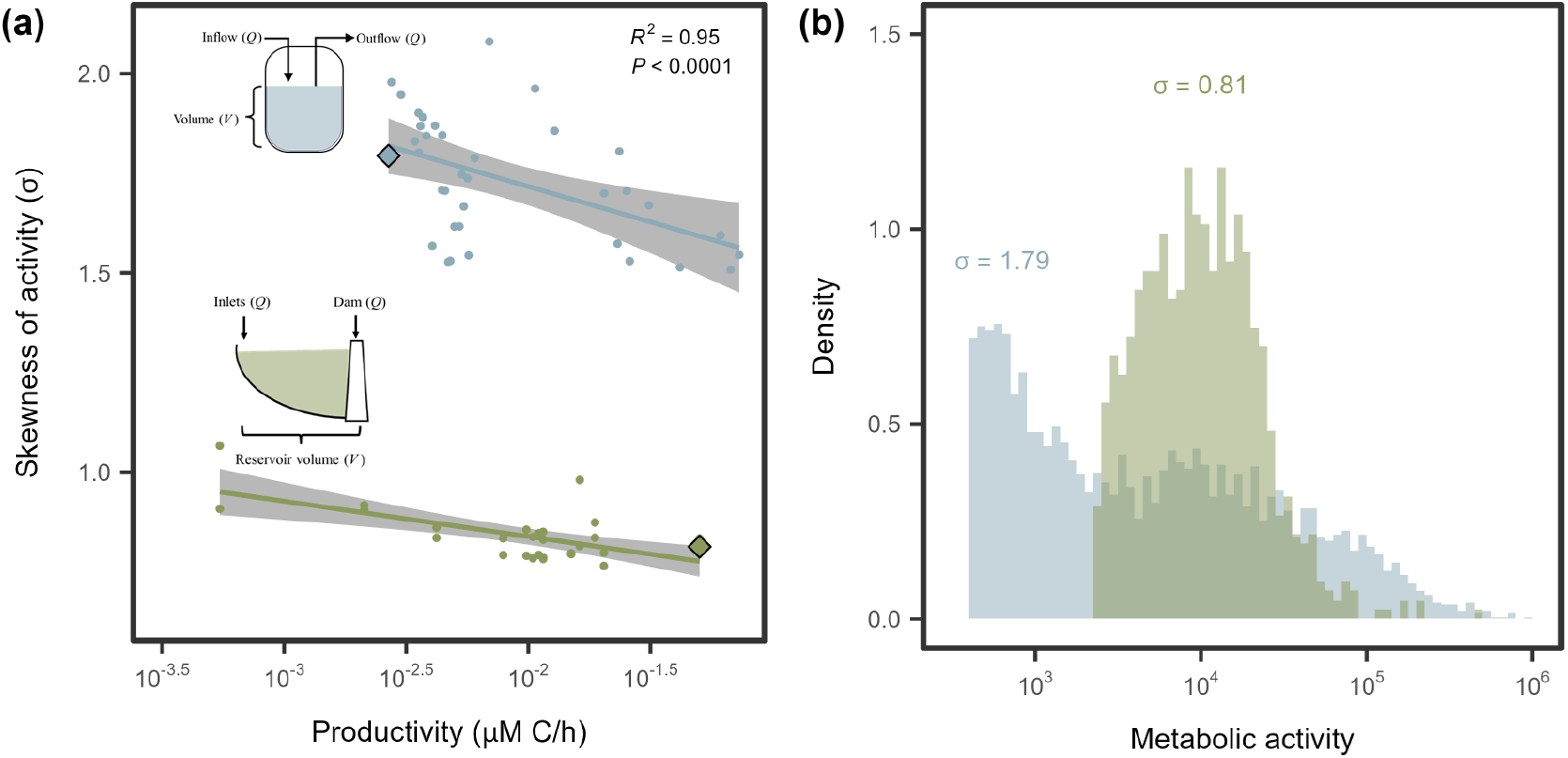
Productivity reduces metabolic inequality. **(A)** In both continuous culture (blue) and environmental samples (green), metabolic activity became less skewed with increasing productivity. Lines and shaded regions represent linear regression fits and 95% confidence intervals, respectively (Table S6). **(B)** Two representative samples, highlighted by diamond-shaped symbols in **(A)**, illustrate contrasts in distribution shape. While both are best fit by the lognormal model, the more skewed sample (blue, *σ* = 1.79) from a continuous culture with a low dilution rate shows a broader range of metabolic activity, with a pronounced tail extending to *∼* 10^6^ RFU. The less skewed sample (green, *σ* = 0.81) from a reservoir site furthest from the dam outlet shows a more even distribution, with values extending only to *∼* 10^5^.

Our findings run counter to the common notion that “the rich get richer and the poor get poorer.” One possible explanation is that low-activity individuals invest more in acquiring resources as availability increases, because the marginal gains are greater. Alternatively, highly active individuals may experience diminishing returns under resource-abundant conditions, particularly if uptake or storage capacity becomes saturated. Our results are also consistent with observations from non-biological systems, such as economic and organizational structures, where increases in resources benefit a larger fraction of the population rather than a select few [37]. Together, these findings suggest that greater resource availability may promote more equitable distributions of output across complex systems, rather than reinforcing existing inequalities.

### 5. Cross-ecosystem consistency in metabolic inequality

To assess the generality of metabolic inequality, we examined its prevalence across diverse ecosystems. In oceans [52–55], lakes (Fig. 4), soils [56], sediments [57, 58], and guts [59], the most active 20% of individuals accounted for 38% to 90% of total metabolic activity, consistent with Pareto-like dynamics (Fig. S9). Single-cell activity distributions were best described by a lognormal distribution (Fig. S10; Table S7 & S8), outperforming alternative models regardless of ecosystem type. Importantly, these results were robust to differences in how cellular metabolism was quantified, including microautoradiography [52, 53, 55], DNA- and amino acid-based click chemistry [54–56], redox indicators [57, 58], and related approaches [59].

We next examined how variation in metabolic inequality is structured across ecosystem types, given that environmental conditions and community composition differ significantly among habitats. Using the intraclass correlation coefficient from a random-intercept model to partition variance in the lognormal skew parameter (*σ*), we found that ecosystem type explained 44% of the total variance (Fig. 6). The datasets analyzed here span ecosystems with wide ranges of redox state, temperature, pressure, solar input, pH, and modes of energy and carbon acquisition. Yet across these distinct habitats, from loamy soils to deep ocean sediments, we observe overlapping ranges of *σ*, indicating that ecosystem-specific signatures do not override broader patterns of metabolic inequality.

**Figure 6.**
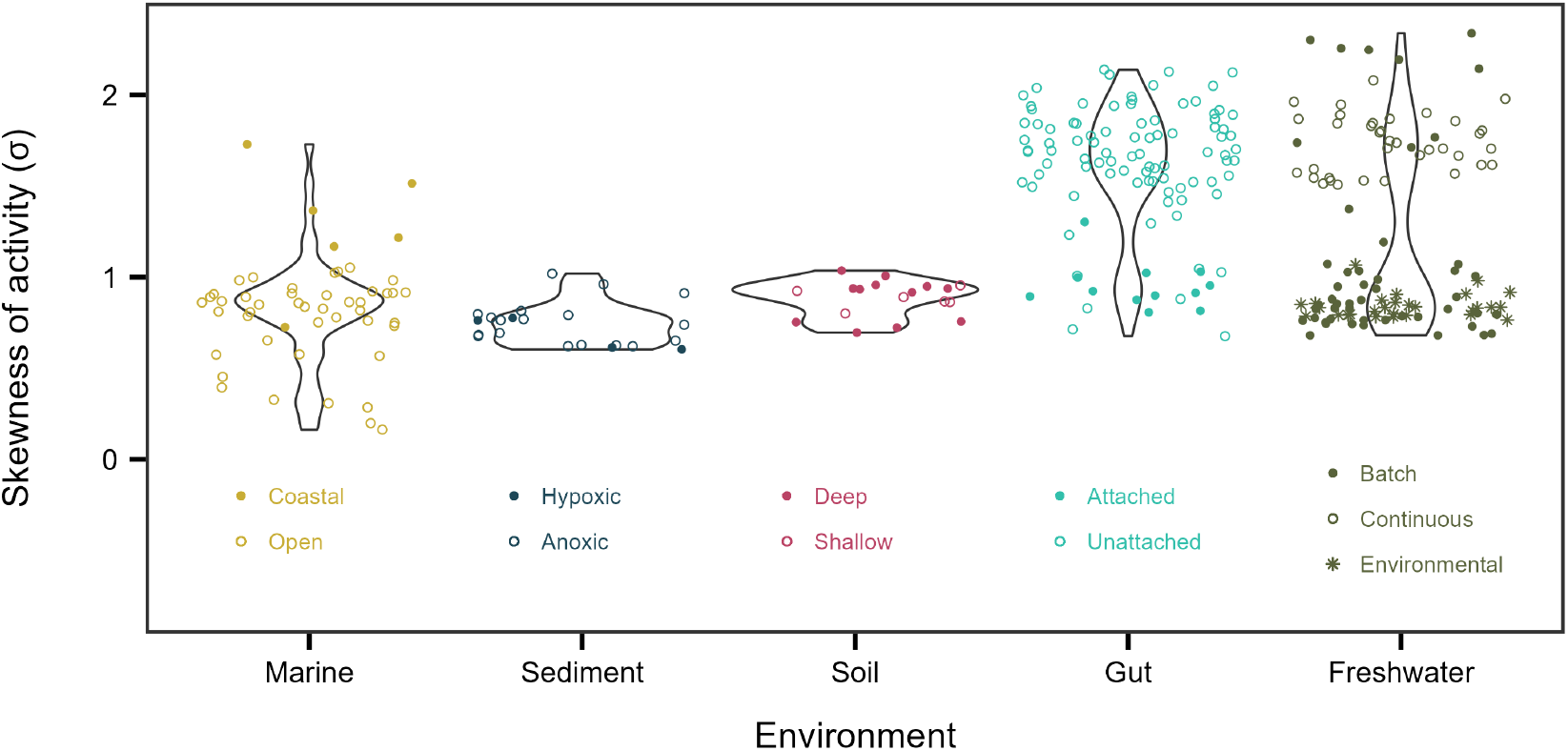
Metabolic inequality across ecosystems. Each point in a violin plot represents the skew in metabolic activity among individuals within a community, measured by the *σ* parameter from lognormal fits.

More than half of the variation in skewness occurred within ecosystems, indicating that metabolic inequality is strongly shaped by habitat-specific conditions. For example, within marine systems, *σ* varied systematically between open-ocean and coastal habitats, while in gut-associated communities it differed between attached and free-living assemblages. These patterns suggest that spatial structure, lifestyle, and productivity (Fig. 5) modulate the magnitude of inequality within ecosystems. In addition, temporal dynamics likely amplify these effects. In our experiments, some of the largest shifts in *σ* reflected expansions and contractions in growth and cell division driven by fluctuations in resource availability (Figs. 2 & 3). Because such dynamics are widespread across natural, engineered, and host-associated systems, they may underlie variation in metabolic inequality.

### 6. Consequences of metabolic inequality for ecosystem functioning

Our results show that metabolic inequality is widespread and follows a consistent, structured pattern across ecosystems (Fig. 6). However, understanding how this physiological variation influences ecosystem processes such as respiration and primary productivity remains poorly understood. Many relationships between traits and performance are inherently nonlinear, such that variation among individuals does not translate proportionally into system-level outcomes. For example, photosynthesis saturates with light, growth plateaus with nutrient supply, and fitness often follows curved trait landscapes. When such nonlinearities are present, the performance predicted from the average trait value differs from the average performance across individuals. This outcome follows from Jensen’s inequality, which states that for nonlinear functions, *f* (𝔼[*x*]) ≠ 𝔼 [*f* (*x*)], where 𝔼 denotes the expectation (mean) of a trait *x* across individuals.

Consequently, model predictions based on mean trait values can be biased. The magnitude of this bias depends on the curvature of the response function and the variance among individuals [60]. It may also be influenced by higher-order features of the trait distribution, such as skewness, which often co-vary with variance. To quantify these effects, we used a stochastic simulation framework to evaluate how the lognormal distribution of single-cell reductase activity (Fig. 4) influences community-level respiration.

Cellular respiration (*R*) was modeled as a saturating function of per-cell reductase activity (*E*) using the canonical Michaelis–Menten equation (Eq. 1). Community respiration was then obtained by aggregating across individuals, allowing us to assess how distributional properties of *E* influence the emergent system-level response. This formulation shows how nonlinear mapping from metabolic activity to respiration produces aggregate responses predicted by Jensen’s inequality.

To evaluate the effects of metabolic inequality on community respiration, we defined an index of respiration bias (*I*_*R*_). This index compares aggregate respiration predicted under a lognormal distribution of reductase activity (*E*) with that predicted under a Gaussian distribution having the same mean. When *I*_*R*_ = 1, respiration is equivalent under Gaussian and lognormal distributions of *E*, indicating no net effect of distributional shape on the aggregated response. Simulations show that respiration bias increases with the skewness (*σ*) of the lognormal distribution and is amplified when the functional response saturates more gradually at higher values of *K*_*m*_ (Fig. 7). Under empirically observed levels of skewness (Figs. 5 & 6), we estimate that community respiration could be overestimated by as much as 60%.

**Figure 7.**
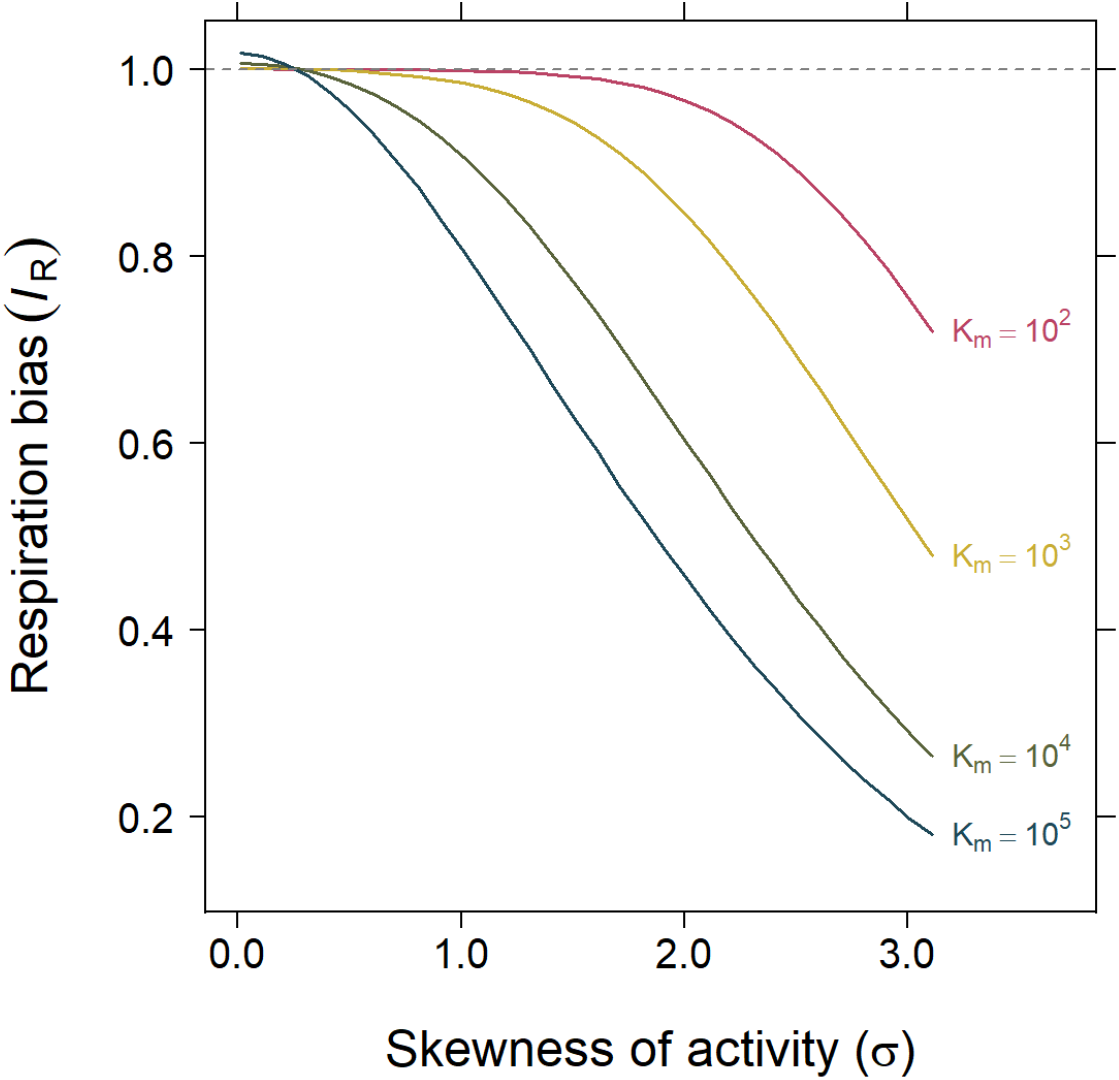
Consequences of metabolic inequality. Simulations show how the index of respiration bias (*I*_*R*_) varies with the skewness (*σ*) of lognormally distributed metabolic activity. *I*_*R*_ is defined as the ratio of aggregate community respiration predicted under a lognormal distribution of reductase activity (*E*) to that predicted under a Gaussian distribution with the same mean. Single-cell respiration (*R*) is modeled using a Michaelis–Menten response to reductase activity (*E*), and community respiration is obtained by aggregating across cells (i.e., summing or averaging individual rates). Each curve corresponds to a different value of the half-saturation constant (*K*_*m*_), which governs the degree of nonlinearity in the functional response. Bias is greatest at high skew and low *K*_*m*_, where the response saturates at lower values of *E*, amplifying the effects of metabolic inequality.

Increasingly, ecosystem and Earth system models incorporate greater biological realism to improve forecasts and better capture feedbacks[26, 61]. For example, some models incorporate microbial traits related to nutrient acquisition, enzymatic activity, thermal sensitivity, and carbon use efficiency [62, 63]. However, most lack explicit representations of metabolic heterogeneity within microbial biomass pools [64, 65]. Instead, they treat individuals as physiologically identical under a common set of environmental conditions. Our results suggest that this simplification overlooks a fundamental feature of microbial communities. Incorporating skewness into even a basic, mechanistic model of microbial respiration reveals that metabolic inequality can substantially alter predicted ecosystem fluxes (Fig. 7). Together, these findings underscore the need to move beyond homogeneous biomass pools and explicitly represent individual-level variation in metabolic activity. Such representation could be achieved through agent-based models that allow distributional structure to emerge dynamically, or by incorporating parameterized activity distributions into trait-based and ecosystem models to evaluate how metabolic inequality propagates through trophic levels and food webs [66, 67]. Extending these approaches to Earth system models would enable assessment of how microbial inequality influences large-scale biogeochemical dynamics and climate–carbon cycle feedbacks.

## Conclusion

Inequality among individuals is a recurring feature of complex systems. In human societies, unequal distributions of wealth influence access to food, healthcare, and education [68]. Although shaped by economic and institutional structures, similar patterns emerge in natural systems. Examples include dominance hierarchies in primates, territoriality in birds, and resource caching in rodents [69]. Together, these cases suggest that inequality is a general property of living systems with consequences for individual fitness and collective function [70].

Here, we extend this perspective to microbial systems, showing that metabolic activity is unequally distributed among cells within complex communities. Activity distributions are consistently right-skewed and well described by a lognormal model, with a minority of cells accounting for a disproportionate share of total function. This pattern is consistent with multiplicative processes operating within large, interacting populations. Although the magnitude of inequality varies with resource availability, habitat structure, and productivity, its widespread occurrence points to an organizing principle of microbial metabolism.

Our findings suggest that metabolic inequality may shape species interactions and community stability. If highly active individuals are more likely to be grazed by predators, infected by viruses, or engage in syntrophic partnerships, then energy and resources may become concentrated within a subset of the population. This concentration may in turn focus ecological risk and interaction strength, with consequences for resilience and coexistence. Recognizing and quantifying these inequalities provides a foundation for linking individual-level variation to community dynamics, ecosystem functioning, and uncovering broader principles that govern complex adaptive systems.

## Methods

### Growth regimes

To investigate how metabolic activity is distributed among individuals, we sampled complex microbial communities under different growth regimes. First, we propagated freshwater samples under batch-culture conditions to determine how the distribution of metabolic activity changes over a growth cycle. We inoculated 1 mL of lake water from Goodman Lake, a meso-eutrophic surface mine lake in Greene-Sullivan State Park, Indiana, USA (39.013 ^*°*^N, 87.236 ^*°*^W) into replicate (*n* = 5) Erlenmeyer flasks (25 mL) containing Lysogeny Broth (LB) medium (4 mL), which were then incubated on a shaker table (200 rpm) at 37 ^*°*^C. At 45-min intervals, we measured biomass via optical density (OD_600_ nm) using a BioPhotometer plus (Eppendorf, Hamburg, Germany) and took samples for flow cytometry as described below. To characterize the growth curve dynamics, we fit optical density (OD_600_ nm) measurements with a logistic growth model using the ‘nls’ function (‘stats’ R package, v. 4.3.2).

Second, to determine how the distribution of metabolic activity changes with community growth rate, we manipulated the dilution rate in a set of continuous flow reactors (*n* = 36). We inoculated each of these chemostats (40 mL) with samples from University Lake, a meso-eutrophic reservoir located in Griffy Woods, Bloomington, Indiana, USA (39.189 ^*°*^N, 86.503 ^*°*^W; [71]). Using a combination of peristaltic pumps and manual pipetting, we delivered twice-autoclaved lake water as growth medium and established dilution rates spanning over seven orders of magnitude (10^*−*6.5^ to 10^0.5^ h). As a source of immigration, we added a dilute suspension of lake sediment daily, corresponding to 1% of the estimated total cell turnover [72]. After 20 d, we destructively harvested each chemostat, measured bacterial productivity (BP) using a tritiated leucine incorporation assay [73], and collected samples for flow cytometry as described below.

Third, to determine the distribution of metabolic activity of microorganisms in a natural ecosystem, we obtained water samples from University Lake, which is a small (6.5 ha) reservoir exhibiting predictable spatial patterns along hydrological flow paths, where gradients in nutrient and light availability shape microbial diversity and productivity [71, 74]. Samples were collected every 25 m from the stream inlet to the dam. Upon return to the laboratory, we measured bacterial productivity (BP) using a tritiated leucine incorporation assay [73] and collected samples for flow cytometry as described below.

### Single-cell metabolic activity

We measured single-cell metabolism using BacLight™RedoxSensor™Green (RSG; excitation/emission: 490/520, Invitrogen), a fluorescent probe that emits stable green fluorescence when reduced by intracellular reductases. These enzymes mediate cellular redox reactions involving electron transfer across a range of metabolic pathways, including but not limited to respiration. Because RSG responds broadly to intracellular redox activity rather than targeting a specific enzyme or pathway, it serves as a general indicator of metabolic activity under both aerobic and anaerobic conditions. Previous studies have shown that RSG fluorescence is positively correlated with bacterial activity and viability across diverse taxa and environments [57, 75, 76]. To stain samples, we diluted each sample 1:100 in phosphate buffered saline (pH = 7.4) and added 1 *µ*L RSG to 1 mL of the diluted sample. We then incubated the samples at 37 °C in complete darkness for 15 min. To preserve fluorescence for analysis, we fixed the cells by adding 25 *µ*L of 25% glutaraldehyde and incubated the samples for an additional 30 min at 25 °C in the dark. Samples were stored at 4 °C until they were run on a NovoCyte 3000 flow cytometer (Agilent, Santa Clara, CA, USA) equipped with a 50 mW laser-emitting light. RSG fluorescence was captured with the 488 nm laser using a 530/30 nm filter. For each sample, we collected 10,000 events with a forward scatter height (FSC-H) greater than 300 (arbitrary units), using a gain of 453 and a flow rate of 14.7 *µ*L min^*−*1^. These settings minimized noise from small FSC and SSC values and ensured that the event rate remained below 1,000 events per second, reducing the likelihood of coincident cell detection.

Once the flow cytometry events were collected, we identified live cells using electronic gating with NovoExpress (NovoCyte software, v. 1.4.1). We first gated for single-cells using a side-scatter height vs. area (SSC-H vs. SSC-A) plot to exclude aggregated cells and instrument noise (Fig. S11A). Next, we removed background signal by applying an RSG fluorescence versus event count gate based on cell-free PBS controls (Fig. S11B). To ensure that viable cells with low RSG fluorescence were not excluded, we validated this gate against unstained live cell samples. Flow cytometry data were then exported as .fcs files and imported into R using the Bioconductor ‘flowCore’ package (v. 2.14.1) for downstream analysis.

### Bacterial isolates and phylogenetic analyses

We isolated bacteria from surface water collected at University Lake (see above) by plating dilutions onto R2A agar and incubating plates at 25 °C in the dark. All strains were purified through multiple transfers of single colonies before assessing single-cell metabolic activity using the staining and flow cytometric procedures described above, after harvesting cells following 24 h of growth in 1/10 R2A broth on a shaker (200 rpm) at 25 °C. We identified each strain by sequencing the 16S rRNA gene. Genomic DNA was extracted using a Microbial DNA Isolation Kit (MoBio), and the 16S rRNA gene was amplified using primers 8F and 1492R following established protocols [77]. Sequences were aligned using MAFFT with a high-accuracy global alignment strategy [78], and a maximum likelihood phylogeny was inferred using RAxML-NG (v1.2) under a GTR+Γ model of sequence evolution, with branch support assessed using 200 bootstrap replicates [79]. Finally, skewness (*σ*) of the lognormal fits was mapped onto the phylogeny using the ape package [80], and phylogenetic signal was assessed using Pagel’s *λ* with the phytools package [81] in R.

### Statistical comparison of model fits

To assess the degree of metabolic inequality in a sample, we quantified the share of total activity contributed by the top 20% most active individuals. We then examined how this measure of skew varied across growth conditions in samples collected from batch cultures, continuous cultures (chemostats), and environmental systems. To characterize the underlying distributions of the metabolic data, we fitted six candidate models: uniform, bimodal, Gaussian, gamma, lognormal, and Pareto. For each sample, we fit the raw metabolic activity values using maximum likelihood estimation with the ‘mle2’ function from the ‘bbmle’ package employing the ‘L-BFGS-B’ method (version 1.0.25). Given that metabolic activity cannot be negative, we used a truncated Gaussian with a lower bound of 0 and no upper bound. The probability distribution function, parameters, starting values, and parameter bounds for each candidate model are provided in Table S9.

We determined the best fitting model for each sample by comparing model performance using a suite of goodness-of-fit metrics. First, we calculated *r*^2^ values for each sample and used ANOVA followed by a *post hoc* Tukey test to determine the best model for each growth condition. Second, we calculated the Kolmogorov-Smirnov statistic (K-S; ‘ks.test’ function) to assess the agreement between the observed data and the distribution predicted by the fitted model. The statistic ranges from 0 (complete agreement) to 1 (no overlap between distributions).

Third, for each sample we selected the best fitting model based on the lowest Akaike Information Criterion (AIC) and Bayesian Information Criterion (BIC) values. Models were considered meaningfully different when the absolute difference in information criterion scores (|ΔIC|) exceeded 2. We visualized model performance using quantile-quantile (Q-Q) plots of the observed metabolic activity against the predicted metabolic activity, subsampled to 10,000 cells per model. Last, to estimate an overall *r*^2^ value for each candidate model across all growth conditions, we randomly subsampled 5,000 cells from the full freshwater dataset (*∼*600,000 cells) across 114 samples. We repeated this 10,000 times, calculating the *r*^2^ for predicted vs. observed values in each iteration and averaged the results to obtain a bootstrapped 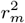 value. We determined the best fitting model for bacterial isolates and samples from across multiple environments as described for freshwater samples. The Gaussian distribution however was not tested on these additional samples as it was shown to be a poor fit for freshwater metabolic activity distributions. The variance in the lognormal skew parameter (*σ*) explained by ecosystem type was calculated as the intraclass correlation coefficient with matrix as the predictor.

### Productivity-inequality relationships

We measured bacterial productivity (BP) by tracking the incorporation of 3H-leucine into microbial communities [73]. We spiked 1.5 mL samples with ^3^H-leucine for a final concentration of 50 nM. After incubating the samples for 1 h at 25 °C, we stopped the reactions by adding 300 *µ*L of 3 mM trichloroacetic acid (TCA). We then washed the cells with 0.3 mM TCA to remove any unincorporated radioactive leucine before covering the samples in scintillation fluid for measurement on a Tri-Carb 2100TR Liquid Scintillation Counter (instrument efficiency: 64.3%; Packard Instrument Company, Meriden, CT, USA). For each sample, we included a voucher replicate in which leucine incorporation was stopped immediately after the isotope spike to determine background radioactivity prior to biological uptake. We converted counts per minute (CPM) to disintegrations per minute (DPM) and used voucher-corrected values to estimate the amount of leucine incorporated. These values were then converted to rates of carbon assimilation (*µ*M C h^*−*1^) using estimates of the fraction of leucine in protein and cellular carbon per protein [83].

To evaluate how productivity influences the distribution of metabolic activity, we examined the relationship between bacterial productivity (BP) and the shape of metabolic activity in the two growth regimes (continuous culture and environmental samples). In the lognormal model, the *σ* parameter (standard deviation in log-space) reflects the degree of skew in metabolic activity. We tested the effect of BP on *σ* model fits using multiple regression with indicator variables [71, 84] where BP was treated as a continuous predictor variable and growth condition (i.e., continuous culture vs. environmental samples) as a categorical variable. An interaction term was included to test whether the relationship between BP and *σ* differed between growth conditions. All statistical analyses were conducted in R (v 4.3.2; R Core Team, 2024).

### Simulation model

We developed a stochastic simulation model to examine how the statistical distribution of enzyme activity levels among individual cells influences community-level respiration. For each simulation, we sampled *n* = 10^6^ cells with independently drawn enzyme levels (*E*_*i*_).

Baseline enzyme activity was drawn from a Gaussian distribution, *E*_*i*_ *∼ N* (*µ, σ*^2^), with *µ* = 10^5^ and *σ* = 0.25*µ*. Negative values arising from the Gaussian distribution were truncated to a small positive constant for numerical stability.

Respiration (*R*_*i*_) for each cell was calculated according to the Michaelis-Menten function,

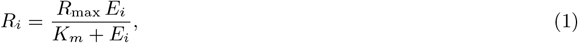

where *R*_max_ = 1 and half-saturation constants (*K*_*m*_) ranged from 10^2^ to 10^5^.

To evaluate the effect of distributional skew, we replaced the Gaussian with lognormal activity distributions, *E*_*i*_ *∼*Lognormal(*µ*_log_, *σ*^2^), where *µ*_log_ is the mean of the log-transformed variable. The lognormal distribution was parameterized such that the arithmetic mean of *E*_*i*_ was held constant at *µ* = 10^5^ while the log-space standard deviation (*σ*) varied from 0.01 to 3.5. This was achieved by solving

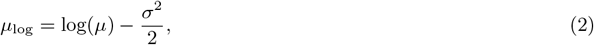

which ensures that 𝔼[*E*_*i*_] = *µ* for all values of *σ*.

For each *K*_*m*_–*σ* combination, we computed mean respiration across all cells,

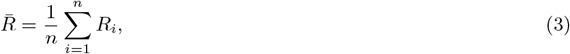

and expressed it as a ratio relative to the Gaussian baseline, which we define as an index of respiration bias,

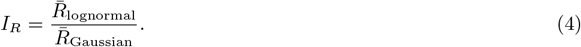

Because total community respiration is the sum across individuals (i.e., proportional to 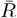 for fixed population size), this ratio also reflects differences in aggregate respiration. Ratios *>* 1 indicate that the lognormal distribution yields higher predicted respiration than the Gaussian baseline, whereas ratios *<* 1 indicate lower respiration. This approach captures the influence of distributional skew (and associated changes in variance) while holding mean activity constant, allowing us to quantify how increasing inequality among cells alters predicted community-level respiration.

## Acknowledgments

We acknowledge support from the US National Science Foundation (DEB-1934554 and DBI-2022049 to JTL), the US Army Research Office Grant (W911NF2210014 to JTL), the National Aeronautics and Space Administration (80NSSC20K0618 to JTL), the Alexander von Humboldt Foundation (to JTL), and the National Institutes of Health (R35GM161372 to JTL). We thank WR Shoemaker, KJ Locey, DR Schoolmaster, LC Moyle, JA Lau, and SR Hall for valuable discussions; S Kang, C Hassel, and BK Lehmkuhl for technical assistance; and C Amano, E Couradeau, D Goudeau, GJ Herndl, DL Kirchman, LM Lindsay, RR Malmstrom, TR Northern, BN Orcutt, E Sintes, SP Smriga for access to and interpretation of published single-cell activity data from different ecosystems. Data and code can be found at http://github.com/LennonLab/MetabolicDistribution.

