## Supplement for "Metabolic inequality in microbial communities"

### 1. Metabolic inequality relationships

We tested the effect of biomass and the three phases of growth (lag, exponential, and stationary) on the share of total activity contributed by the top 20% most active individuals in batch culture using generalized additive models (GAMs). GAMs are linear models where the response variable depends on smoothed functions of predictor variables, allowing for non-linear responses. We fit GAMs to our data using the 'gam' function (R package mgcv, version 1-9.1) with thin plate regression splines as the basis function (s()) using restricted maximum likelihood (REML) to estimate the smoothing parameter ($\lambda$) while not restricting the dimensions of the basis used for the smoothing term ($k$) ([1]). We report the results of a modified Wald test if the coefficient of the smoothed terms is equal to zero ($p$-values, $\alpha = 0.5$) as well as deviance explained for each GAM ([2]). We selected models based on a conditional Akaike information criterion (AIC) where GAM fit AIC method corrects for effective degrees of freedom (edf) of the model.

**Table S1. Goodness-of-fit for candidate model of metabolic activity.** Summary statistics for $r^2$ and Kolmogorov-Smirnov statistics for the four tested distributions (lognormal, gamma, Gaussian, and Pareto). Estimated marginal means and standard error (EM means ± SE) are shown for each distribution within each growth condition. Each growth condition has a degree of freedom (df) and each distribution has a 95% confidence interval (95% CI) and a letter rank (Tukey rank) generated from a *post hoc* Tukey test. In all cases, "Gaussian" refers to a truncated Gaussian distribution, with bounds (0, ∞].

| | Distribution | EM means ± SE | 95% CI | Tukey rank |
|---|---|---|---|---|
| | | $r^2$ | | |
| Batch Culture df = 190 | Lognormal | 0.897 ± 0.013 | (0.870, 0.923) | a |
| | Gamma | 0.919 ± 0.013 | (0.893, 0.946) | a |
| | Gaussian | 0.923 ± 0.014 | (0.895, 0.952) | a |
| | Pareto | 0.777 ± 0.013 | (0.751, 0.804) | b |
| Continuous Culture df = 137 | Lognormal | 0.988 ± 0.009 | (0.969, 1.006) | a |
| | Gamma | 0.936 ± 0.009 | (0.917, 0.954) | b |
| | Gaussian | 0.516 ± 0.010 | (0.497, 0.535) | c |
| | Pareto | 0.986 ± 0.009 | (0.968, 1.004) | a |
| Environmental sampling df = 88 | Lognormal | 0.990 ± 0.010 | (0.971, 1.009) | a |
| | Gamma | 0.950 ± 0.010 | (0.931, 0.969) | b |
| | Gaussian | 0.619 ± 0.018 | (0.583, 0.655) | c |
| | Pareto | 0.990 ± 0.010 | (0.971, 1.009) | a |
| | | Kolmogorov-Smirnov statistic | | |
| Batch Culture df = 190 | Lognormal | 0.208 ± 0.013 | (0.181, 0.234) | b |
| | Gamma | 0.173 ± 0.013 | (0.146, 0.199) | b |
| | Gaussian | 0.184 ± 0.014 | (0.156, 0.212) | b |
| | Pareto | 0.341 ± 0.013 | (0.315, 0.368) | a |
| Continuous Culture df = 137 | Lognormal | 0.072 ± 0.007 | (0.0.058, 0.086) | c |
| | Gamma | 0.161 ± 0.007 | (0.147, 0.174) | b |
| | Gaussian | 0.637 ± 0.007 | (0.622, 0.651) | a |
| | Pareto | 0.090 ± 0.007 | (0.076, 0.104) | c |
| Environmental sampling df = 88 | Lognormal | 0.068 ± 0.010 | (0.049, 0.087) | d |
| | Gamma | 0.139 ± 0.010 | (0.120, 0.158) | c |
| | Gaussian | 0.550 ± 0.018 | (0.515, 0.585) | a |
| | Pareto | 0.276 ± 0.010 | (0.257, 0.295) | b |

**Table S2. *Post hoc* Tukey test pairwise contrasts of distribution fit $r^2$ values and Kolmogorov-Smirnov statistics within each growth condition.** Each contrast (Contrast) is given along with the estimate (Estimate) and standard error (SE) of the contrast. The $t$-ratio statistic ($t$-ratio) is given along with the $p$-value ($p$-value) for each contrast.

| | Contrast | Estimate | SE | $t$-ratio | $p$-value |
|---|---|---|---|---|---|
| | | $r^2$ | | | |
| Batch culture df = 190 | lognormal – gamma | -0.023 | 0.019 | -1.19 | 0.6371 |
| | lognormal – gaussian | -0.027 | 0.020 | -1.36 | 0.5234 |
| | lognormal – pareto | 0.119 | 0.019 | 6.28 | <0.0001 |
| | gamma – gaussian | -0.004 | 0.020 | -0.22 | 0.9963 |
| | gamma – pareto | 0.142 | 0.019 | 7.47 | <0.0001 |
| | gaussian – pareto | 0.146 | 0.020 | 7.44 | <0.0001 |
| Continuous culture df = 137 | lognormal – gamma | 0.052 | 0.013 | 3.96 | 0.0007 |
| | lognormal – gaussian | 0.472 | 0.013 | 35.14 | <0.0001 |
| | lognormal – pareto | 0.002 | 0.013 | 0.14 | 0.9991 |
| | gamma – gaussian | 0.420 | 0.013 | 31.27 | <0.0001 |
| | gamma – pareto | -0.050 | 0.013 | -3.82 | 0.0011 |
| | gaussian – pareto | -0.470 | 0.013 | -35.00 | <0.0001 |
| Environmental sampling df = 88 | lognormal – gamma | 0.040 | 0.014 | 2.93 | 0.0218 |
| | lognormal – gaussian | 0.371 | 0.020 | 18.18 | <0.0001 |
| | lognormal – pareto | 0.000 | 0.014 | 0.02 | 1.0000 |
| | gamma – gaussian | 0.331 | 0.020 | 16.22 | <0.0001 |
| | gamma – pareto | -0.040 | 0.014 | -2.91 | 0.0233 |
| | gaussian – pareto | -0.371 | 0.020 | -18.16 | <0.0001 |
| | | Kolmogorov-Smirnov statistic | | | |
| Batch culture df = 190 | lognormal – gamma | 0.035 | 0.019 | 1.84 | 0.2576 |
| | lognormal – gaussian | 0.024 | 0.020 | 1.21 | 0.6193 |
| | lognormal – pareto | -0.133 | 0.019 | -7.02 | <0.0001 |
| | gamma – gaussian | -0.011 | 0.020 | -0.57 | 0.9415 |
| | gamma – pareto | -0.168 | 0.019 | -8.86 | <0.0001 |
| | gaussian – pareto | -0.157 | 0.020 | -8.01 | <0.0001 |
| Continuous culture df = 137 | lognormal – gamma | -0.088 | 0.010 | -8.91 | <0.0001 |
| | lognormal – gaussian | -0.565 | 0.010 | -55.67 | <0.0001 |
| | lognormal – pareto | -0.018 | 0.010 | -1.80 | 0.2788 |
| | gamma – gaussian | -0.476 | 0.010 | -46.95 | <0.0001 |
| | gamma – pareto | 0.071 | 0.010 | 7.12 | <0.0001 |
| | gaussian – pareto | 0.547 | 0.010 | 53.91 | <0.0001 |
| Environmental sampling df = 88 | lognormal – gamma | -0.071 | 0.013 | -5.29 | <0.0001 |
| | lognormal – gaussian | -0.482 | 0.020 | -23.92 | <0.0001 |
| | lognormal – pareto | -0.208 | 0.013 | -15.51 | <0.0001 |
| | gamma – gaussian | -0.411 | 0.020 | -20.39 | <0.0001 |
| | gamma – pareto | -0.137 | 0.013 | -10.22 | <0.0001 |
| | gaussian – pareto | 0.274 | 0.020 | 13.57 | <0.0001 |

**Table S3. Goodness-of-fit for candidate model of bacterial isolate metabolic activity.** Summary statistics for $r^2$ and Kolmogorov-Smirnov statistics for the three tested distributions. Estimated marginal means and standard error (EM means ± SE) are shown for each distribution within each growth condition. Each growth condition has a degree of freedom (df) and each distribution has a 95% confidence interval (95% CI) and a letter rank (Tukey rank) generated from a *post hoc* Tukey test.

| | Distribution | EM means ± SE | 95% CI | Tukey rank |
|---|---|---|---|---|
| $r^2$ | | | | |
| Isolates df = 66 | Lognormal | 0.967 ± 0.011 | (0.945, 0.989) | a |
| | Gamma | 0.951 ± 0.011 | (0.929, 0.973) | a |
| | Pareto | 0.879 ± 0.011 | (0.857, 0.902) | b |
| Kolmogorov-Smirnov statistic | | | | |
| Isolates df = 66 | Lognormal | 0.095 ± 0.013 | (0.069, 0.121) | b |
| | Gamma | 0.123 ± 0.013 | (0.097, 0.148) | b |
| | Pareto | 0.374 ± 0.013 | (0.348, 0.399) | a |

**Table S4. *Post hoc* Tukey test pairwise contrasts of distribution fit $r^2$ values and Kolmogorov-Smirnov statistics for bacterial isolates.** Each contrast (Contrast) is given along with the estimate (Estimate) and standard error (SE) of the contrast. The $t$-ratio statistic ($t$-ratio) is given along with the $p$-value ($p$-value) for each contrast.

| | Contrast | Estimate | SE | $t$-ratio | $p$-value |
|---|---|---|---|---|---|
| | | | $r^2$ | | |
| Isolates df = 66 | lognormal – gamma | 0.016 | 0.016 | 1.01 | 0.5763 |
| | lognormal – pareto | 0.087 | 0.016 | 5.56 | <0.0001 |
| | gamma – pareto | 0.072 | 0.016 | 4.56 | <0.0001 |
| | | Kolmogorov-Smirnov statistic | | | |
| Isolates df = 66 | lognormal – gamma | -0.027 | 0.018 | -1.50 | 0.2990 |
| | lognormal – pareto | -0.279 | 0.018 | -15.20 | <0.0001 |
| | gamma – pareto | -0.251 | 0.018 | -13.70 | <0.0001 |

**Table S5. Output from generalized additive model for supplemental figures.** Response variables for each generalized additive model (Fit model) are shown with deviance explained and $P$-value for the total model (Model $p$-value). Each smoothed term (Term) has an effective degree of freedom (edf) and $F$-value ($F$-statistic). Significance of each smoothed term (Term) was determined from term $p$-values ($p$-value) ($\alpha$ = 0.05). Restricted maximum likelihood values (REML) for each model are shown.

| Fit Model | Deviance Explained | Term | Edf | $F$-statistic | REML | $p$-value |
|---|---|---|---|---|---|---|
| Activity of top 20% cells | 0.96 | s(Biomass) | 3.98 | 103.4 | -9.82 | $< 0.0001$ |
| $log_{10}$(Bacterial productivity) | 0.92 | s(Distance from Dam) | 7.44 | 55.70 | -58.2 | $< 0.0001$ |
| $log_{10}$(Bacterial productivity) | 0.92 | s(Growth rate) | 4.98 | 58.13 | -12.40 | $< 0.0001$ |
| Skewness of activity ($\sigma$) | 0.28 | s(Growth rate) | 2.83 | 2.74 | -14.01 | 0.0457 |

**Table S6. Summary statistics for relationship between metabolic activity distributions and bacterial productivity.** Output from indicator variables multiple regression model testing how the skew of metabolic activity (lognormal $\sigma$) is altered by bacterial productivity (BP). Model coefficients (Estimate) are shown for each term (Term) of the model. Significant linear relationships were determined from model $p$-values ($\alpha = 0.05$). Test statistics ($t$-Statistic) and standard errors (SE) for each model term are also shown.

| Response | Term | Estimate | SE | $t$-Statistic | $p$-value |
|---|---|---|---|---|---|
| Lognormal $\sigma$ | Intercept | 1.361 | 0.090 | 15.14 | $< 0.0001$ |
| | $Log_{10}$(BP) | -0.178 | 0.042 | -4.22 | $< 0.0001$ |
| | Environmental samples | -0.700 | 0.130 | -5.39 | $< 0.0001$ |
| | $Log_{10}$(BP) $\times$ Environmental samples | 0.089 | 0.062 | 1.45 | 0.153 |

**Table S7. Goodness-of-fit for candidate model of cross-environment metabolic activity.** Summary statistics for $r^2$ and Kolmogorov-Smirnov statistics for the three tested distributions. Estimated marginal means and standard error (EM means ± SE) are shown for each distribution within each growth condition. Each growth condition has a degree of freedom (df) and each distribution has a 95% confidence interval (95% CI) and a letter rank (Tukey rank) generated from a *post hoc* Tukey test.

| | Distribution | EM means ± SE | 95% CI | Tukey rank |
|---|---|---|---|---|
| | | $r^2$ | | |
| Cross-environment df = 546 | Lognormal | 0.967 ± 0.003 | (0.961, 0.974) | a |
| | Gamma | 0.957 ± 0.003 | (0.950, 0.963) | b |
| | Pareto | 0.960 ± 0.003 | (0.954, 0.967) | ab |
| | | Kolmogorov-Smirnov statistic | | |
| Cross-environment df = 546 | Lognormal | 0.101 ± 0.004 | (0.093, 0.109) | b |
| | Gamma | 0.106 ± 0.004 | (0.097, 0.114) | b |
| | Pareto | 0.161 ± 0.004 | (0.153, 0.169) | a |

**Table S8. *Post hoc* Tukey test pairwise contrasts of distribution fit $r^2$ values and Kolmogorov-Smirnov statistic across environments.** Each contrast (Contrast) is given along with the estimate (Estimate) and standard error (SE) of the contrast. The $t$-ratio statistic ($t$-ratio) is given along with the $p$-value ($p$-value) for each contrast.

| | Contrast | Estimate | SE | $t$-ratio | $p$-value |
|---|---|---|---|---|---|
| $r^2$ | | | | | |
| Cross-environment df = 546 | lognormal – gamma | 0.011 | 0.005 | 2.40 | 0.0437 |
| | lognormal – pareto | 0.007 | 0.005 | 1.56 | 0.2624 |
| | gamma – pareto | -0.004 | 0.005 | -0.84 | 0.6784 |
| Kolmogorov-Smirnov statistic | | | | | |
| Cross-environment df = 546 | lognormal – gamma | -0.004 | 0.006 | -0.76 | 0.7273 |
| | lognormal – pareto | -0.060 | 0.006 | -10.18 | <0.0001 |
| | gamma – pareto | -0.055 | 0.006 | -9.42 | <0.0001 |

**Table S9. Probability distribution functions and starting values for parameters estimated.** We fit the resulting metabolic activity distribution of each sample to four distributions, the lognormal, gamma, truncated Gaussian, and Pareto distribution (Distribution). Here we show the probability distribution function (Equation) for each distribution as well as the parameters estimated for each (Parameter). Starting values given to the 'mle2' function using the method 'L-BFGS-B' are given (Starting value). In some cases, the starting value of a parameter was dependent on the sample values, and this is indicated by 'median(sample)' in the table.

| Distribution | Equation | Parameter | Starting value |
|---|---|---|---|
| Lognormal | $PDF = \frac{1}{x\sigma\sqrt{2\pi}}\ exp\left(-\frac{(ln\ x-\mu)}{2\sigma^2}\right)$ | Log mean ($\mu$) | 1 |
| | | Log std deviation ($\sigma$) | median(sample) |
| Gamma | $PDF = \frac{1}{\Gamma(\alpha)\beta^\alpha} x^{\alpha-1} e^{-\frac{x}{\beta}}$ where $\Gamma(\alpha) = (\alpha - 1)!$ | Shape ($\alpha$) | 1 |
| | | Scale ($\beta$) | 10 |
| Truncated Gaussian | $PDF = \begin{cases} 0, x \leq 0 \\ \frac{1}{\sigma} \frac{\varphi(\frac{x-\mu}{\sigma})}{1-\phi(\frac{-\mu}{\sigma})}, x > 0 \end{cases}$ where $\varphi(x) = \frac{1}{\sqrt{2\pi}} e^{-\frac{1}{2}x^2}$ and $\phi(x) = \frac{1}{2\pi}\int_{-\infty}^{x} e^{-t^2/2} dt$ | Mean ($\mu$) | median(sample) |
| | | Std deviation ($\sigma$) | 100 |
| Pareto | $PDF = \frac{\alpha\ x_m^\alpha}{x^{\alpha+1}}$ | Shape ($\alpha$) | 1 |
| | | Log std deviation ($\chi_m$) | median(sample) |
| Uniform | $PDF = \begin{cases} \frac{1}{b-a},\ a \leq x \leq b \\ 0,\ x < a\ or\ x > b \end{cases}$ | Minimum ($a$) | 1 |
| | | Maximum ($b$) | median(sample) |

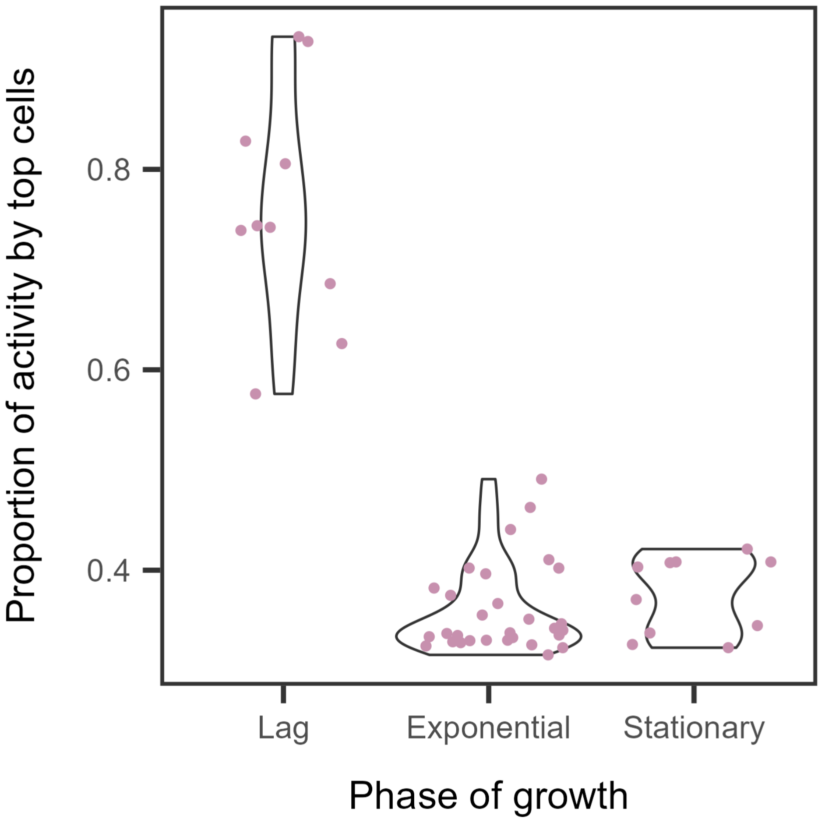


**Figure S1. In batch culture, the proportion of activity performed by the top 20% of individuals is dependent on the phase of growth.** The percentage of total activity performed by the top 20% of individuals in a community is highest in lag phase. In exponential phase, the proportion of activity performed by the top 20% of individuals decreases, with most samples having 30% of activity performed by these individuals. In stationary phase the proportion remains low, but the distribution of the values shifts upward, increasing as resources are consumed.

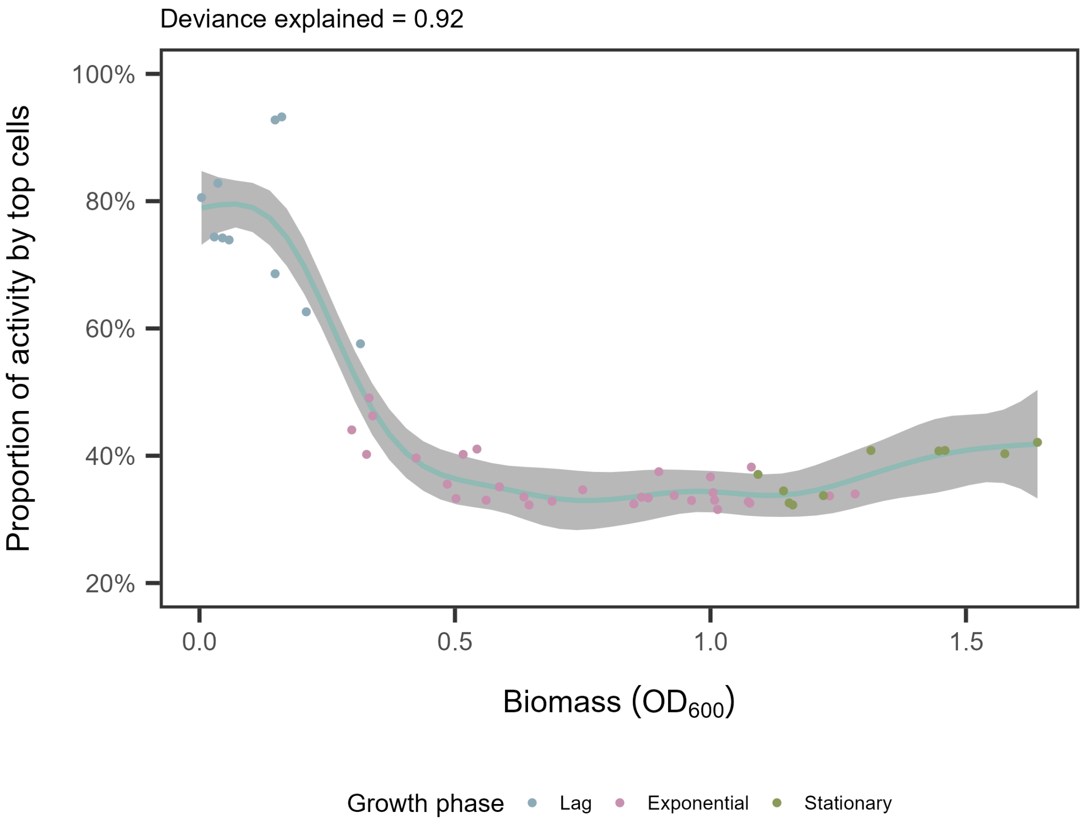


**Figure S2. Proportion of activity by the most active cells (top 20%) is dependent on community biomass.** The percentage of total activity performed by the top 20% of individuals in a community decreases quickly during lag phase as cells adjust to the new environment. In exponential phase, the activity performed by the top 20% of individual cells is the lowest as this is where growth rates are expected to be the most uniform. As resources are exhausted, the communities enter stationary phase and again show an increase in the percentage of total activity performed by the top 20% of individual cells. Lines and shading represent fits and 95% CIs of GAM regressions, respectively. The percentage of deviance explained by the model is reported above each plot.

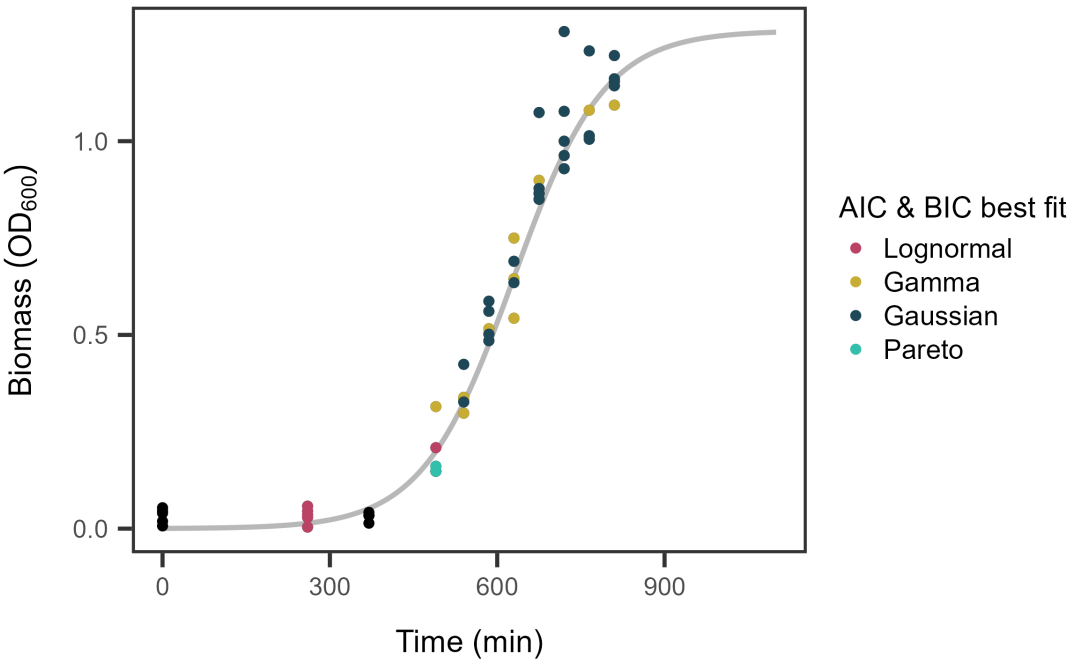


**Figure S3. Best fit distribution from Akaike and Bayesian information criterion (AIC and BIC respectively) varies along a growth curve.** Biomass for five replicate lake communities was measured for a growth curve along with the metabolic activity of each replicate community at each time point. The metabolic activity distribution for each sample at each time point was fit with a lognormal, gamma, Gaussian, and Pareto distribution and AIC and BIC values were calculated for each. Points are colored based on the best fit distribution for each sample as determined by lowest AIC and BIC value with a $|\Delta \text{ IC}|$ greater than 2 as best fit model calls based on AIC and BIC were the same for all sample. Samples where $OD_{600\ nm}$ was measured but flow cytometry was not performed are shown in black.

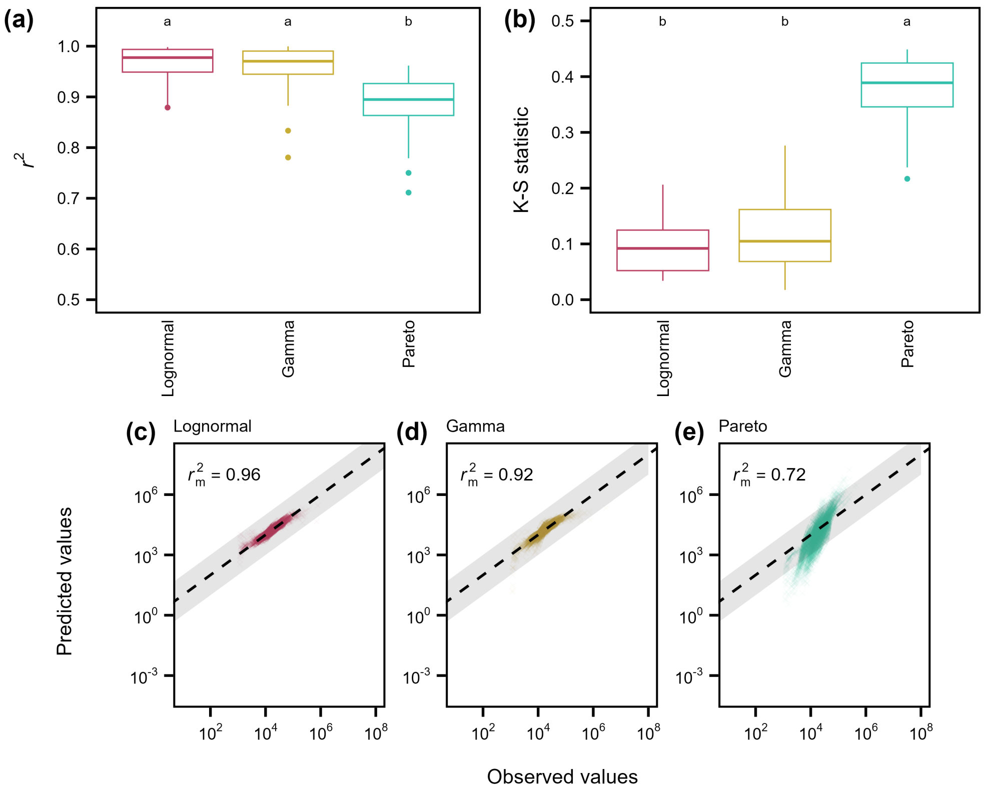


**Figure S4. Metabolic activity of bacterial isolates follows a lognormal distribution.** Metabolic activity distributions from bacterial isolates were fit with three candidate models: lognormal, gamma, and Pareto distributions. (a) We calculated the $r^2$ value for each sample's fit to each candidate model. Across all additional environments, the lognormal distribution performed comparably to the Pareto distribution but outperformed the gamma distribution. (b) Kolmogorov-Smirnov (K-S) statistics, where lower values indicate better fit, showed that the lognormal distribution performed comparably to the gamma distribution but outperformed the Pareto distribution. Letters rankings were assigned based on *post hoc* Tukey tests. Predicted values based on the model fits were plotted against the observed values of metabolic activity for the (c) lognormal ($r_m^2$=0.99), (D) gamma ($r_m^2$=0.97), and (E) Pareto ($r_m^2$=0.96). In each quantile-quantile plot, observed and predicted metabolic activity values are shown for 10,000 sub-sampled individuals from the bacterial isolates. Solid lines represent the 1:1 line (perfect agreement) and shaded regions represent ± 1 order of magnitude in activity.

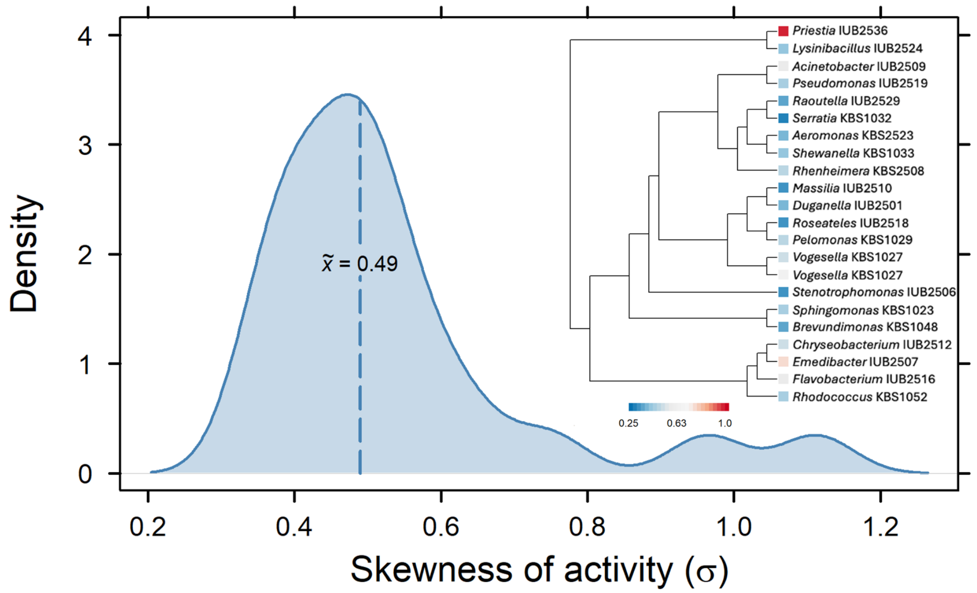


**Figure S5. Skewness of bacterial isolate metabolic distributions.** The mean skewness of the 22 isolates from University Lake is 0.49, while the range of observed skewness values was 0.35-1.1. A phylogeny generated from the 16S rRNA gene sequences of the isolates is inset in the figure, with the color of the phylogeny tips indicating the skewness of metabolic activity ($\sigma$)

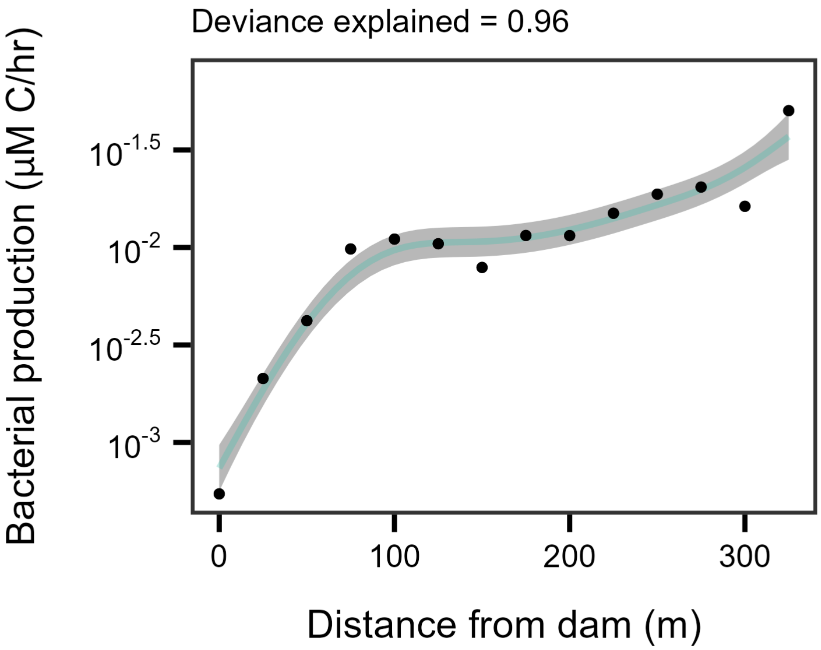


**Figure S6. Bacterial productivity along hydrological flowpath.** Productivity is highest near the three stream inlets, decreasing more sharply in the 100 m closest to the dam. Lines and shading represent fits and 95% CIs of GAM regressions, respectively. The percentage of deviance explained by the model is reported above each plot

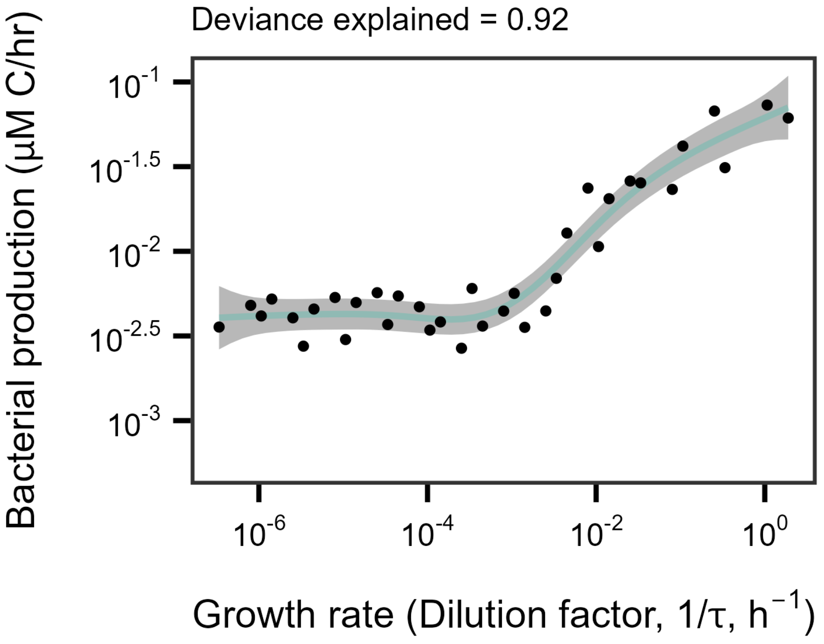


**Figure S7. Bacterial productivity as function of growth rate in chemostats.** Growth rate is estimated to be equal to the dilution factor ($1/\tau$) of a chemostat. Lines and shading represent fits and 95% CIs of GAM regressions, respectively. The percentage of deviance explained by the model is reported above each plot.

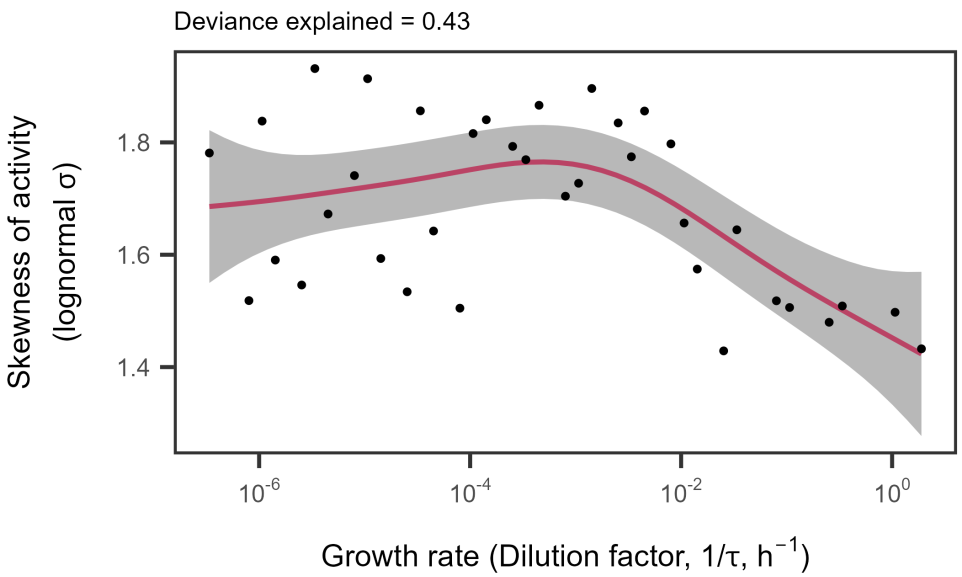


**Figure S8. Skewness of metabolic activity ($\sigma$) as function of growth rate in chemostats.** Growth rate is estimated to be equal to the dilution factor ($1/\tau$) of a chemostat. Lines and shading represent fits and 95% CIs of GAM regressions, respectively. The percentage of deviance explained by the model is reported above each plot.

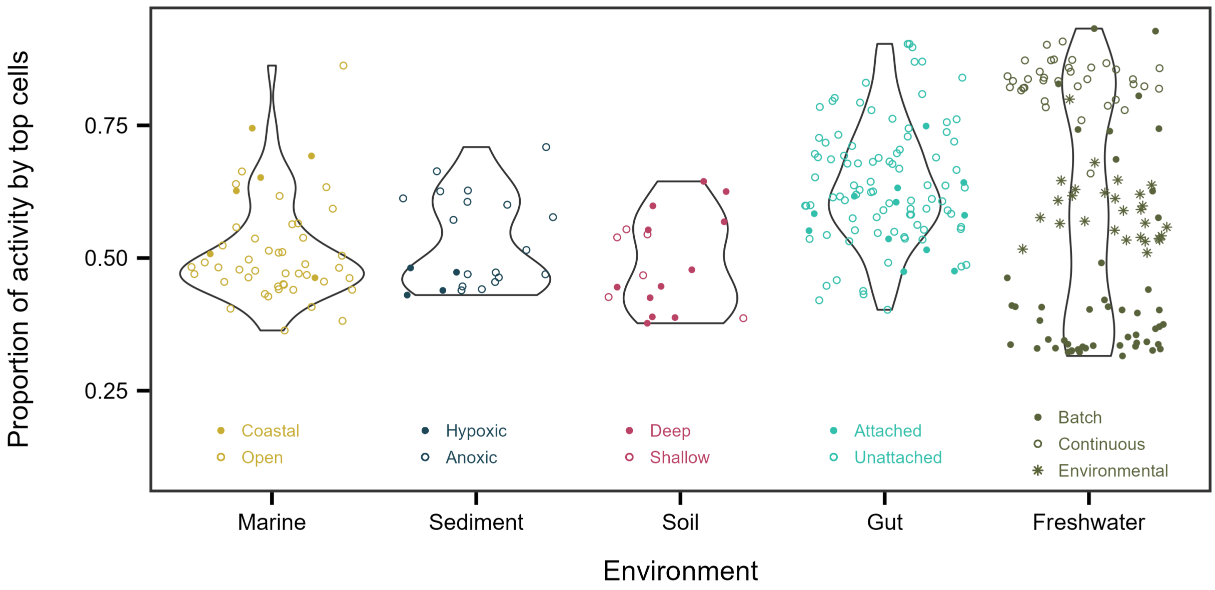


**Figure S9. The proportion of activity performed by the top 20% most active individuals varied both within and between environments.** Among the four additional environments, marine, sediment, soil, and gut, and the freshwater samples previously described, the proportion of variance among environments (0.04) is much smaller than the proportion of variance within environments (0.96).

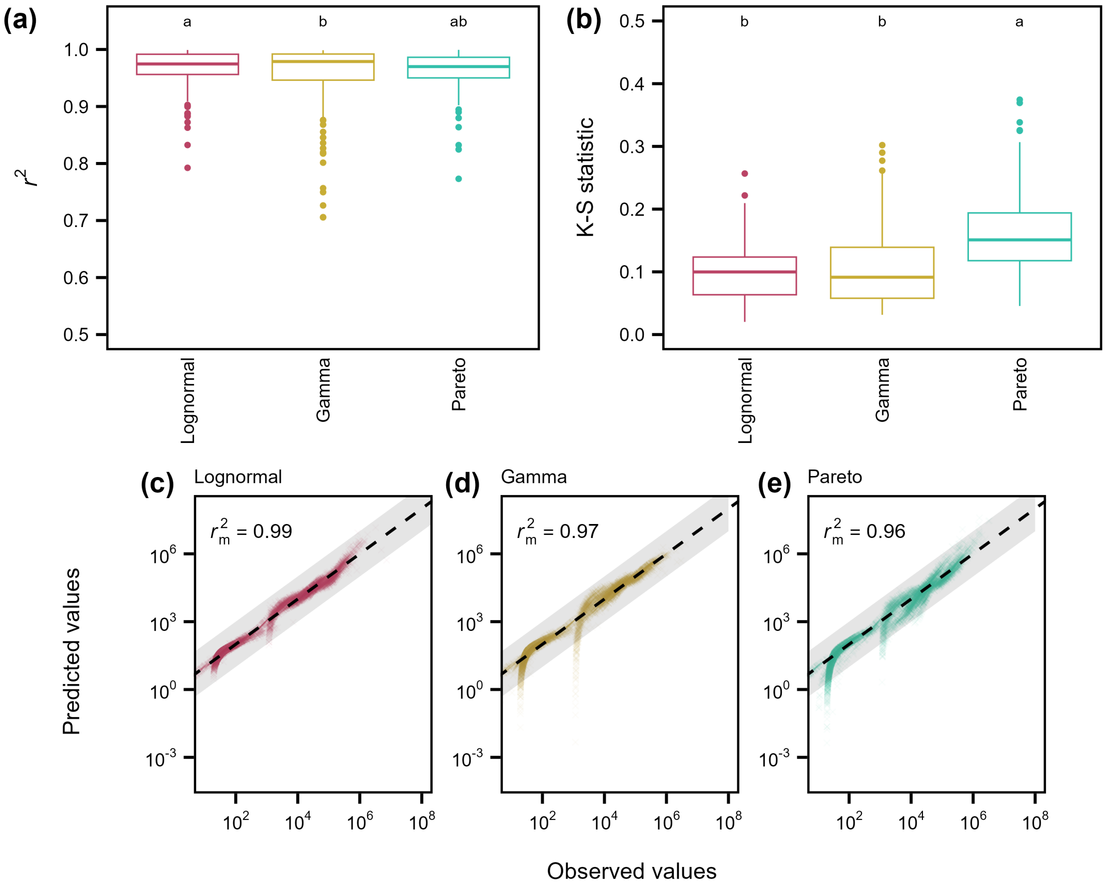


**Figure S10. Across ecosystems, metabolic activity follows a lognormal distribution.** Metabolic activity distributions from all samples across four additional environments (soil, sediment, marine, and gut) were fit with three candidate models: lognormal, gamma, and Pareto distributions. (a) We calculated the $r^2$ value for each sample's fit to each candidate model. Across all additional environments, the lognormal distribution performed comparably to the Pareto distribution but outperformed the gamma distribution. (b) Kolmogorov-Smirnov (K-S) statistics, where lower values indicate better fit, showed that the lognormal distribution performed comparably to the gamma distribution but outperformed the Pareto distribution. Letters rankings were assigned based on *post hoc* Tukey tests. Predicted values based on the model fits were plotted against the observed values of metabolic activity for the (c) lognormal ($r_m^2$=0.99), (D) gamma ($r_m^2$=0.97), and (E) Pareto ($r_m^2$=0.96). In each quantile-quantile plot, observed and predicted metabolic activity values are shown for 10,000 sub-sampled individuals from the additional environments. Solid lines represent the 1:1 line (perfect agreement) and shaded regions represent ± 1 order of magnitude in activity.

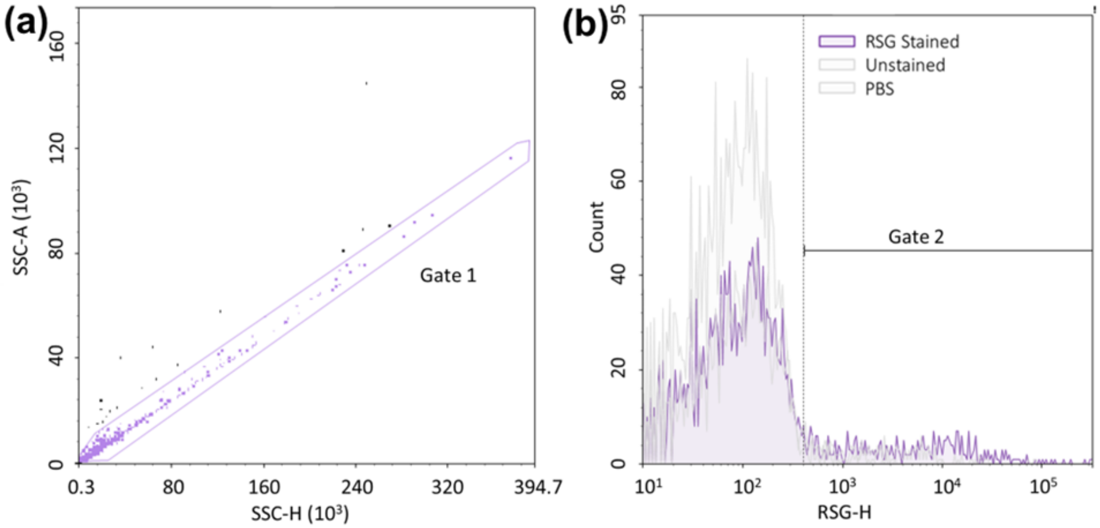


**Figure S11. Flow cytometry gating for metabolic activity.** (a) Cells were gated on a side scatter height vs. side scatter area (SSC-H vs. SSC-A) plot for single-celled organisms to remove aggregates and a portion of baseline cytometer noise from the sample. (b) On a RedoxSensor Green (RSG) fluorescence count plot, cells were gated for RSG activity. This gate was positioned to remove machine noise captured with unstained phosphate buffered saline (PBS) samples while avoiding removal of cells that were found in the unstained sample, meaning that live and active cells, regardless of activity level, were captured with this gate.
